# Reduction of protein disulfide isomerase results in open conformations and stimulates dynamic exchange between structural ensembles

**DOI:** 10.1101/2020.12.07.414680

**Authors:** Mathivanan Chinnaraj, Robert Flaumenhaft, Nicola Pozzi

## Abstract

Protein disulfide isomerase (PDI) is a ubiquitous redox-regulated enzyme that interacts with hundreds of client proteins intracellularly and extracellularly. It comprises two redox-sensitive domains, each hosting the conserved catalytic motif CxxC, two redox-insensitive protein-binding domains, and three linkers. Snapshots of oxidized and reduced PDI have been obtained by X-ray crystallography. Yet, how PDI’s structure dynamically changes in response to the redox microenvironment and ligand binding remain unknown. Here, we used multiparameter confocal single-molecule Förster resonance energy transfer (smFRET) and multiple FRET pairs to track the movements of the two catalytic domains with high temporal resolution. Our studies document that, at equilibrium, PDI visits three structurally distinct conformational ensembles, two “open” (O_1_ and O_2_) and one “closed” (C). We show that the redox environment dictates the time spent in each ensemble and the rate at which they exchange. While oxidized PDI samples O_1_, O_2_ and C more evenly and in a slower fashion, reduced PDI predominantly populates O_1_ and O_2,_ and exchanges between them more rapidly, on the sub-millisecond timescale. These findings were not expected based on crystallographic data. Using mutational analyses, we further demonstrate that the two active sites are structurally nonequivalent and that ligands targeting the active sites of reduced PDI shift the equilibrium towards closed conformations of the enzyme. This work introduces a new structural framework that challenges current views of PDI dynamics, helps rationalize the multifaced role of PDI in biology and may assist drug development.

## Introduction

Protein disulfide isomerase (PDI) is an archetypal oxidoreductase responsible for oxidative protein folding in eukaryotes *(1, 2)*. It interacts with hundreds of client proteins catalyzing the formation, rupture, and isomerization of disulfide bonds *(1, 3)*. Since disulfide bonds are essential for achieving tertiary and quaternary structures of proteins but also have important functional roles, its enzymatic activity is essential for life.

PDI comprises 508 amino acids organized in four thioredoxin domains arranged in the order **a, b, b’** and **a**’, followed by an acidic C-terminal tail **(Figure 1A)** *(1, 3)*. Domains **a** (res 18-133) and **a’** (res 137-232) contain the conserved catalytic motif CxxC, whereas the **b** (res 235-348) and **b’** (res 370-479) domains are redox-insensitive; they participate in substrate and cofactor recruitment, not in catalysis. Three linkers connect the four thioredoxin domains.

**Figure 1.**
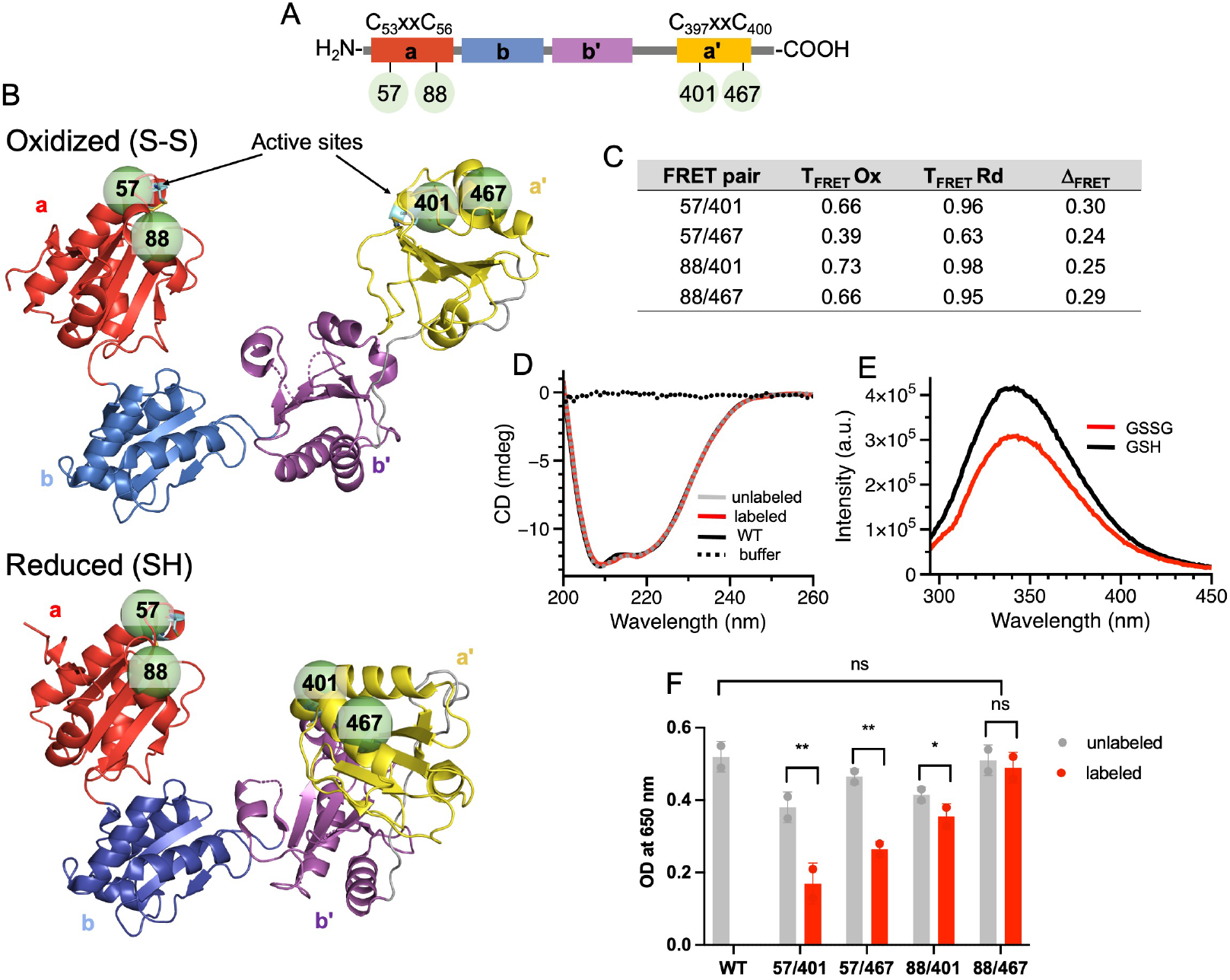
Design, structural and functional integrity of labeled proteins. **A)** Domain structure of human PDI. PDI comprises 508 AA organized in four domains, **a** (red), **b** (blue), **b’** (magenta) and **a’** (yellow), connected by linkers (gray). Domains **a** and **a’** contain the active sites CxxC. **B)** Top view of the X-ray crystal structures of oxidized (S-S, top panel, 4el1) and reduced (SH, bottom panel, 4ekz) PDI documenting a U-shape architecture and movement of the **a’** domain (yellow) toward the **b’** domain (magenta) upon reduction of the active sites. Shown as dotted lines are the distances between the active sites. The green spheres represent residues K57, S88, K401 and K467 selected for smFRET studies. **C)** Theoretical values of energy transfer for oxidized (T_FRET_Ox) and reduced PDI (T_FRET_Rd) estimated with the software FPS after attaching Atto-550 and Atto-467N dyes (R_0_=65Å) at the specified positions. The difference T_FRET_Ox-T_FRET_Rd is shown as Δ_FRET_. **D)** Far-UV CD of PDI 88/467 before (gray) and after (red) labeling compared to PDI wild-type (WT, black). **E)** Reactivity of PDI 88/467 towards oxidized (GSSG, red) and reduced (GSH, black) glutathione probed by intrinsic fluoresce spectroscopy. **F)** Insulin assay testing the reductase activity of proteins before (gray) and after (red) labeling. PDI wild type (WT) served as a control. Each reaction was continuously monitored for 60 min at 650 nm in duplicate. The intensity at 650 nm after 40 min was used as readout of catalytic efficiency. Progress curves are shown in **Figure S1. (**\*\* p<0.001), * p<0.01, n.s. not significant).

While mostly located in the endoplasmic reticulum (ER) *(1)*, PDI can also be found outside the cells, where it plays key regulatory roles in several enzymatic cascades, most notably in the coagulation cascade *(4-6)*. However, in contrast to intracellular PDI, which primarily works as a foldase *(1, 7)*, extracellular PDI mostly works as a reductase or oxidase *(4, 8)*. That is, PDI facilitates the rupture or formation of specific types of disulfide bonds in target proteins, known as allosteric disulfide bonds *(9)*, thus modulating their function by inducing local or global structural rearrangements. Relevant examples of allosteric disulfide bonds regulated by PDI are the ones in the coagulation activator tissue factor *(10)*, in the Antiphospholipid Syndrome autoantigen β_2_GPI *(11)*, and in the membrane receptors αIIbβ3 *(12)* and GpIbα *(13)*, which control platelet activation and aggregability.

Because of the motif CxxC, the catalytic activity of PDI is critically regulated by the microenvironment via cysteine modifications, i.e., oxidation (S-S) or reduction (-SH) *(1)*. In the ER, there are several systems controlling the redox balance so that PDI is mostly oxidized *(14)*. However, in the circulation, such systems are not readily available. Consequently, PDI’s redox state can vary quite significantly. Hence, understanding how the structure of PDI responds to changes in the redox milieu is particularly important for the field of thrombosis and hemostasis as this knowledge could help to define the mechanistic basis of its extracellular function as well as to design compounds capable of targeting redox-specific activities of extracellular PDI for safe anticoagulation.

Over the past two decades, structural, computational, and biophysical studies have provided solid evidence for PDI’s flexibility by documenting large-scale redox-dependent and redox-independent movements of the two catalytic domains *(15-21)*. This led to the hypothesis that PDI operates as a dynamic clamp, which is capable of opening and closing in response to different stimuli. Testing this structure-based hypothesis, however, has been challenging due to several technical limitations. Ensemble methods, although easily accessible, are difficult to interpret on a structural basis since the signal is averaged over multiple conformations, preventing direct identification of distinct functional states. Multidimensional NMR experiments are greatly complicated by the relatively large size of PDI. Finally, X-ray crystallography and cryo-electron microscopy, while providing very detailed information, offer a limited number of structural snapshots. X-ray crystallography may also impose constraints on structural variability.

Recently, pioneering single-molecule studies of PDI have started to emerge in the literature. Most notably, Okumura et al. applied High-Speed Atomic Force Microscopy (HS-AFM) to study PDI conformational dynamics after tethering the protein onto mica sheets *(20)*. In the meantime, our group developed a method to incorporate unnatural amino acids into PDI *(22)*, thus opening the door for single-molecule Förster resonance energy transfer (smFRET) investigations of PDI in solution. In this study, we provide a detailed characterization of the conformational dynamics of PDI in solution using multiparameter confocal smFRET. We chose this setup over other single-molecule approaches, such as total internal reflection, because it enables identification and quantification of large-scale dynamics with high temporal resolution (ns-ms) while minimizing the probability of structural perturbations caused by immobilization of proteins onto a surface *(23-25)*. Our work reveals unanticipated movements of the **a** and **a’** catalytic domains in response to the redox environment and ligand binding and offers a new structural framework that helps rationalize the multifaced role of PDI in biology and may assist drug development efforts.

## Results

### Experimental design

To perform smFRET experiments, donor and acceptor fluorophores must be site-specifically introduced into the protein of interest without perturbing its structure and, ideally, its biological function. Positions 57 and 88 in the **a** domain and 401 and 467 in the **a’** domain were selected based on the currently available X-ray structural data of oxidized **(Figure 1B, top panel)** and reduced **(Figure 1B, bottom panel)** PDI *(16)* to obtain four combinations of labeling positions, two linear (i.e., 57/401 and 88/467) and two diagonal (i.e., 57/467 and 88/401). Residues K57 and K401 are located one position downstream of the active site cysteines C56 and C400 in the **a** and **a’** domain. Residues S88 and K467 are located on the opposite side of the catalytic domains. These four FRET pairs were designed to measure large-scale hinge bending movements of the **a** and **a’** domains reported by the structures and to follow the positioning of the active sites relative to each other. The four FRET pairs were also designed such as we expect significantly higher values of energy transfer (Δ_FRET_>0.2) while transitioning from the oxidized (lower FRET) to the reduced state (higher FRET), as reported in **Figure 1C**.

### Site-specific labeling of catalytically active PDI

Given that PDI’s active sites contain four cysteine residues, donor (Atto550) and acceptor (Atto647N) fluorophores were introduced at the desired positions using biorthogonal chemistry by following a procedure recently developed and validated in our laboratory *(22)*. After purification, doubly labeled PDI 57/401, PDI 57/467, PDI 88/401 and 88/467 were properly folded, as documented by far-UV CD **(Figure 1D** and **Figure S1)** and responded well to redox stimulation, as probed by tryptophan fluorescence spectroscopy **(Figure 1E** and **Figure S1)**. Similar spectral variations induced by GSSG and GSH were found in previous studies for PDI wild-type *(26)*. Importantly, doubly labeled PDI variants remained active when tested for their ability to reduce insulin to a degree consistent with what was expected based on the positioning of the dyes **(Figure 1G** and **Figure S1)**. In fact, the loss of enzymatic activity for doubly labeled PDI 57/401 (67%), PDI 57/467 (49%) and PDI 88/401 (32%) compared to unlabeled proteins, PDI wild-type and doubly labeled PDI 88/467 was anticipated and likely arises from the proximity of residue 57 and residue 401 to the active site cysteines 56 and 400, which interferes with substrate processing.

To rule out position-dependent interactions of the dyes with the protein, we measured steady-state anisotropy and quantum yield for singly labeled donor and acceptor PDI molecules. The values measured for the eight variants are reported in **Table 1**. They are consistent with freely rotating dyes attached to PDI.

**Table 1.**
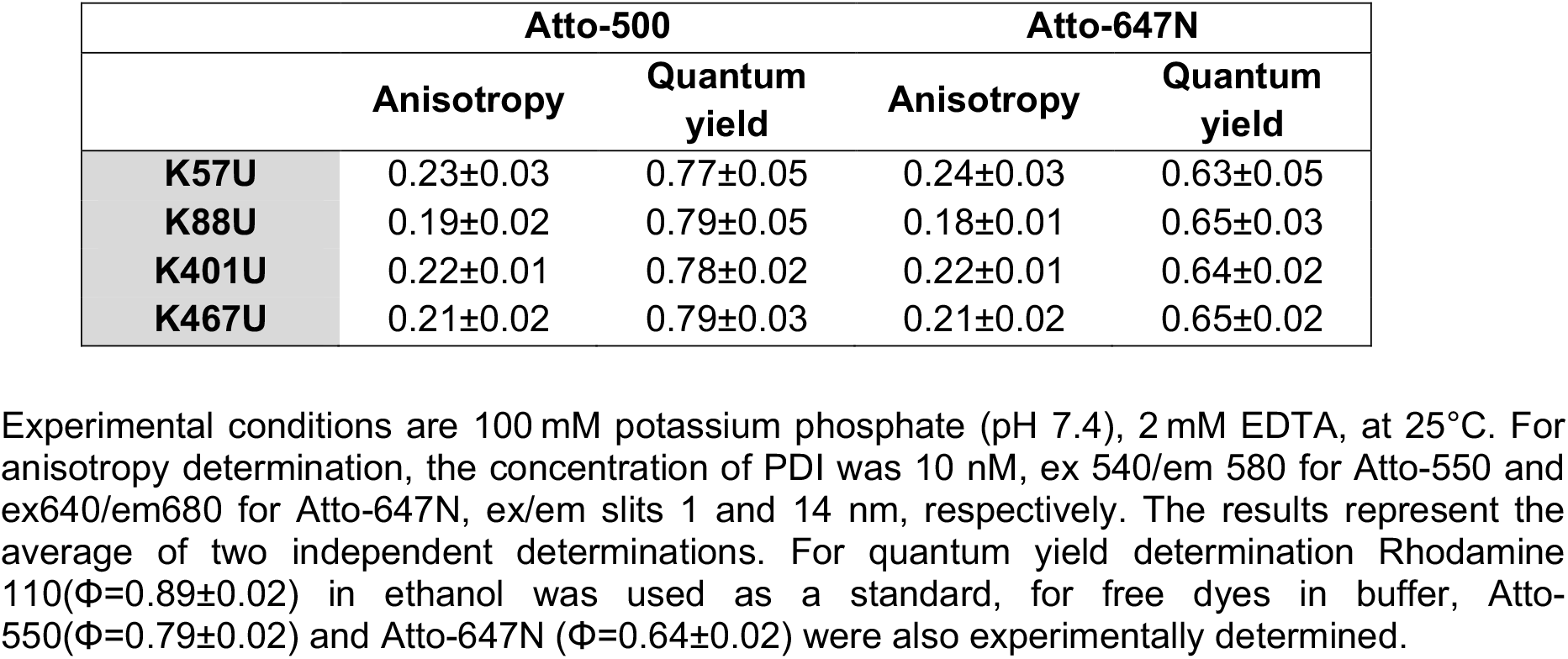
Anisotropy and quantum yield (Φ) values for singly labeled PDI mutants.

|  | Atto-500 |  | Atto-647N |  |
| --- | --- | --- | --- | --- |
|  | Anisotropy | Quantum yield | Anisotropy | Quantum yield |
| **K57U** | 0.23±0.03 | 0.77±0.05 | 0.24±0.03 | 0.63±0.05 |
| **K88U** | 0.19±0.02 | 0.79±0.05 | 0.18±0.01 | 0.65±0.03 |
| **K401U** | 0.22±0.01 | 0.78±0.02 | 0.22±0.01 | 0.64±0.02 |
| **K467U** | 0.21±0.02 | 0.79±0.03 | 0.21±0.02 | 0.65±0.02 |

Experimental conditions are 100 mM potassium phosphate (pH 7.4), 2 mM EDTA, at 25°C. For anisotropy determination, the concentration of PDI was 10 nM, ex 540/em 580 for Atto-550 and ex640/em680 for Atto-647N, ex/em slits 1 and 14 nm, respectively. The results represent the average of two independent determinations. For quantum yield determination Rhodamine 110( $\Phi$ =0.89 ± 0.02) in ethanol was used as a standard, for free dyes in buffer, Atto-550( $\Phi$ =0.79 ± 0.02) and Atto-647N ( $\Phi$ =0.64 ± 0.02) were also experimentally determined.

Taken together, these results indicate that, after purification, fluorescently labeled PDI molecules are properly folded and catalytically active, thus suitable for smFRET studies.

### Conformational dynamics of oxidized and reduced PDI monitored by smFRET

smFRET studies of PDI were performed using a confocal microscope equipped with pulse interleaved excitation, as detailed in the experimental section. This methodology, in addition to enabling simultaneous acquisition of fluorescence intensity and fluorescence lifetime, facilitates isolation and, therefore, analysis of molecules containing the proper donor:acceptor ratio (stoichiometry, S=0.5) while discarding donor only (S=1) and acceptor only (S=0) species, which are irrelevant for our goal **(Figure S2)**. Given that R_0_ of the FRET couple Atto-550/Atto-647N is ∼65Å, interprobe distances from ∼45Å (E=0.9) to ∼90Å (E=0.10) are measured in this study.

The 1D FRET efficiency plots of PDI 57/401, PDI 57/467, PDI 88/401 and PDI 88/467 obtained after size exclusion chromatography (SEC) purification, and therefore under conditions in which the active site cysteines are oxidized *(22)*, are reported in the top panels of **Figure 2A, Figure 2B, Figure 2C** and **Figure 2D**, respectively. The corresponding 2D plots of FRET efficiency vs Stoichiometry (S) are reported in **Figure S3** of the supplementary materials.

**Figure 2.**
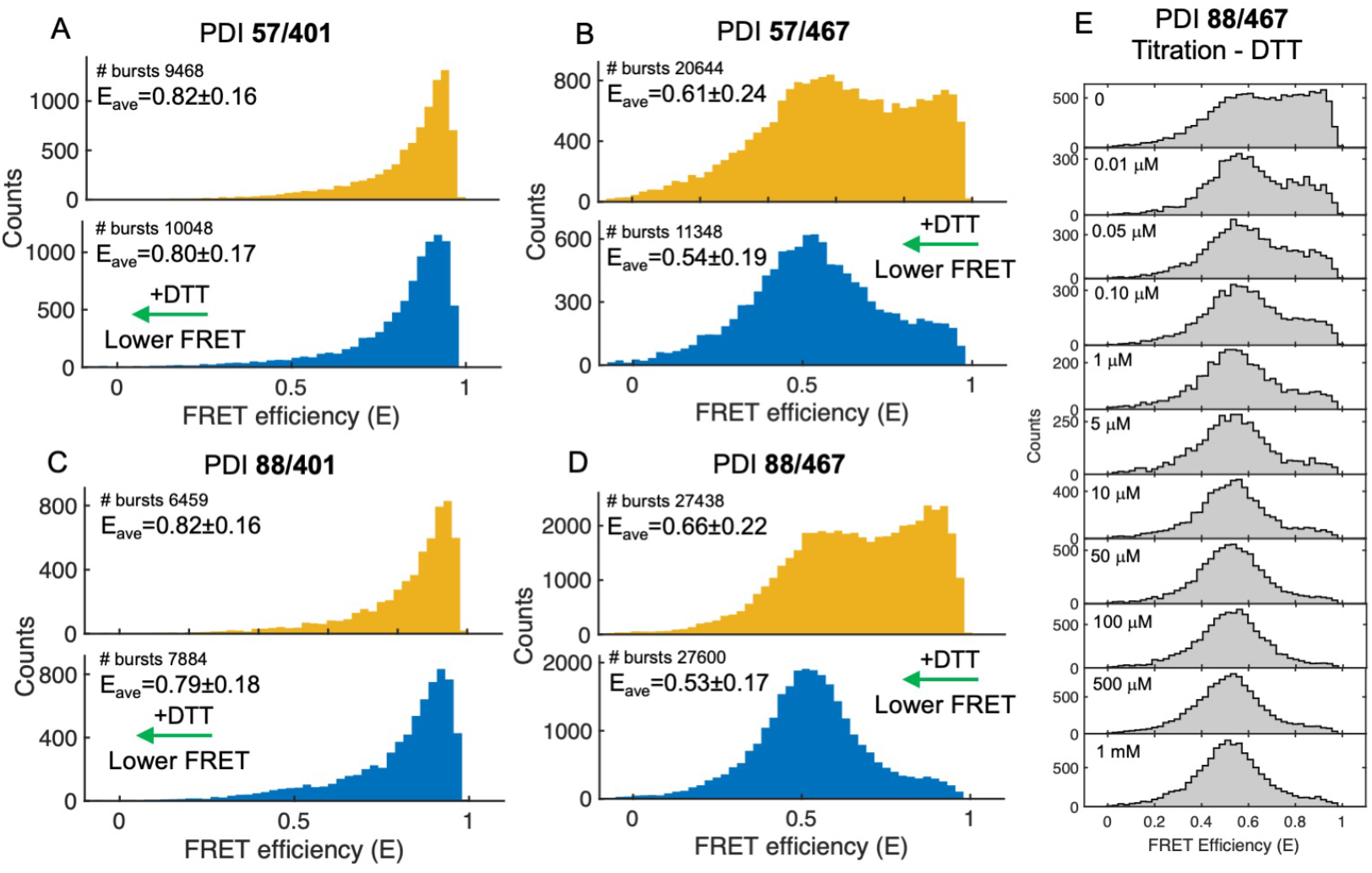
smFRET studies of oxidized and reduced PDI. 1D FRET efficiency histograms of PDI 57/401 **(A)**, PDI 57/467 **(B)**, PDI 88/401 **(C)** and PDI 88/467 **(D)** obtained in the absence (yellow, top panel) and presence (cyan, bottom panel) of 1 mM DTT in Tris 20 mM pH 7.4, 150 mM NaCl, 2 mM EDTA, 0.003% Tween 20. PDI’s concentration was 50-100 pM. Collection time was ∼40 minutes per sample. Molecules with 0.25<S>0.75 were selected. Shown are the number of bursts and mean FRET (E_ave_) ± STDEV. Note how, in the presence of DTT, the signal shifts towards lower FRET (green arrow). **(E)** 1D FRET efficiency histograms of PDI 88/467 collected at increasing concentrations of DTT (0-1 mM). To ensure equilibrium, samples were measured after incubating PDI 88/467 and DTT for 40 minutes at room temperature (20°C).

Oxidized PDI 57/401 and PDI 88/401 were found to adopt a unimodal distribution, which was skewed toward high FRET. For both variants, the mean FRET (E_ave,ox_) was 0.82±0.16. In contrast, oxidized PDI 57/467 and PDI 88/467 displayed broader FRET distributions, spanning almost the entire FRET range. This resulted in lower FRET values (E_ave,ox_=0.61 for PDI 57/467 and E_ave,ox_=0.66 for PDI 88/467) and larger standard deviations (0.24 for PDI 57/467 and 0.22 for PDI 88/467). The differences between these variants are clearly visible by inspecting the plots in **Figure 2**. The values of E_ave,ox_ and S are summarized in **Table 2**.

**Table 2.** Summary of FRET (E) and Stoichiometry values (S) for the PDI variants measured in the absence (top) and presence (bottom) of DTT.

|  | Average<br>E | STD<br>E | Average<br>S | STD<br>S |
| --- | --- | --- | --- | --- |
| **57/401** | 0.82 | 0.16 | 0.48 | 0.094 |
| **57/467** | 0.61 | 0.24 | 0.50 | 0.104 |
| **88/401** | 0.82 | 0.16 | 0.50 | 0.096 |
| **88-467** |  |  |  |  |
| **WT** | 0.66 | 0.22 | 0.52 | 0.098 |
| **AAAA** | 0.64 | 0.22 | 0.51 | 0.099 |
| **AACC** | 0.65 | 0.22 | 0.53 | 0.098 |
| **CCAA** | 0.67 | 0.21 | 0.51 | 0.099 |
| **57/401** | 0.80 | 0.17 | 0.48 | 0.085 |
| **57/467** | 0.54 | 0.19 | 0.51 | 0.100 |
| **88/401** | 0.79 | 0.18 | 0.50 | 0.095 |
| **88-467** |  |  |  |  |
| **WT** | 0.53 | 0.17 | 0.52 | 0.090 |
| **AAAA** | 0.64 | 0.22 | 0.51 | 0.098 |
| **AACC** | 0.56 | 0.21 | 0.50 | 0.090 |
| **CCAA** | 0.54 | 0.16 | 0.51 | 0.085 |

Average FRET and Stoichiometry values and corresponding errors were determined using the BurstBrowser module in PAM.

To transform oxidized PDI into reduced PDI we added the reducing agent dithiothreitol (DTT). When 1 mM DTT was added to oxidized PDI, the FRET signal shifted towards lower values for all the FRET pairs (**Figure 2A, Figure 2B, Figure 2C** and **Figure 2D, bottom panels**, and **Table 2)**. The greatest effect was seen for PDI 88/467 (ΔE_ave,(ox-rd)_=0.13) followed by PDI 57/467 (ΔE_ave,(ox-rd)_=0.07). Smaller variations were measured for PDI 88/401 (ΔE_ave,(ox-rd)_=0.03) and PDI 57/401 (ΔE_ave,(ox-rd)_=0.02). Moreover, PDI 57/467 **(Figure 2B)** and PDI 88/467 **(Figure 2D)** displayed more homogeneous distributions characterized by a significantly smaller standard deviation compared to the oxidized form. The changes observed in the presence of DTT were neither dye nor DTT specific since similar results were obtained by labeling PDI 88/467 with a different combination of dyes and by using GSH as an alternative source of reducing equivalents **(Figure S4)**. Moreover, the effect of DTT on PDI 88/467 was dose-dependent and saturable, as shown in **Figure 2E**. Finally, the changes induced by DTT were also consistent with our previous results obtained with the FRET pair 42/467 *(22)*, which reports an even more pronounced high to low FRET transition in the presence of DTT (ΔE_ave,(ox-rd)_=0.31). We conclude that addition of DTT triggers a profound structural reorganization forcing the catalytic domains to move away from each other. Interestingly, the structural effect elicited on PDI by DTT were qualitatively different from what has been previously reported for the thiol-enzyme quiescin-sulfhydryl oxidase (QSOX) using smFRET *(27)*, implying different sensing mechanisms between families of oxidoreductases.

### Oxidized and reduced PDI undergo rapid conformational dynamics

Data in **Figure 2** document significant differences between the FRET pairs. They also provide preliminary evidence that multiple conformations of oxidized PDI exist at equilibrium, and that PDI 57/467 and PDI 88/467, but not PDI 57/401 nor PDI 88/401, are capable of efficiently visualizing them when labeled with the FRET pair Atto550/647N.

By taking advantage of the way photons are collected and stored in our experiments (i.e., time-correlated single photon counting or TCSPC), we constructed 2D plots in which the transfer efficiency of oxidized and reduced PDI was graphed versus the fluorescence lifetime of the donor in the presence of the acceptor (τ_D(A)_) of each molecule. In these plots, as demonstrated elsewhere *(23–25, 28–30)*, FRET populations that represent conformational states (or ensembles) that either do not exchange or exchange at a rate ∼10 times slower or ∼10 times faster than the molecules’ diffusion time lie on the so-called “static” FRET line, which is the line that describes the theoretical relationship between the values of τ_D(A)_ and the values of energy transfer. By contrast, FRET populations that represent conformational states undergoing dynamic exchange during the observation time deviate from the “static” FRET line and lie on the “dynamic” FRET line, which connects two exchanging states. Because of the very significant effect induced by DTT **(Figure 2D)** and pristine catalytic activity **(Figure 1F)**, we selected PDI 88/467 for our in-depth biophysical analyses. However, similar considerations are applicable and remain valid for the other FRET variants, whose results are reported and briefly discussed in **Figure S5** of the supplementary materials.

For both oxidized **(Figure 3A)** and reduced PDI 88/467 **(Figure 3B)**, we observed two main populations of molecules, connected by a bridge. These two populations were characterized by fluorescence lifetime values centered at ∼0.25 ns and ∼1.59 ns, corresponding to high-FRET and medium-FRET. For simplicity, we called these two populations closed and open, respectively.

**Figure 3.**
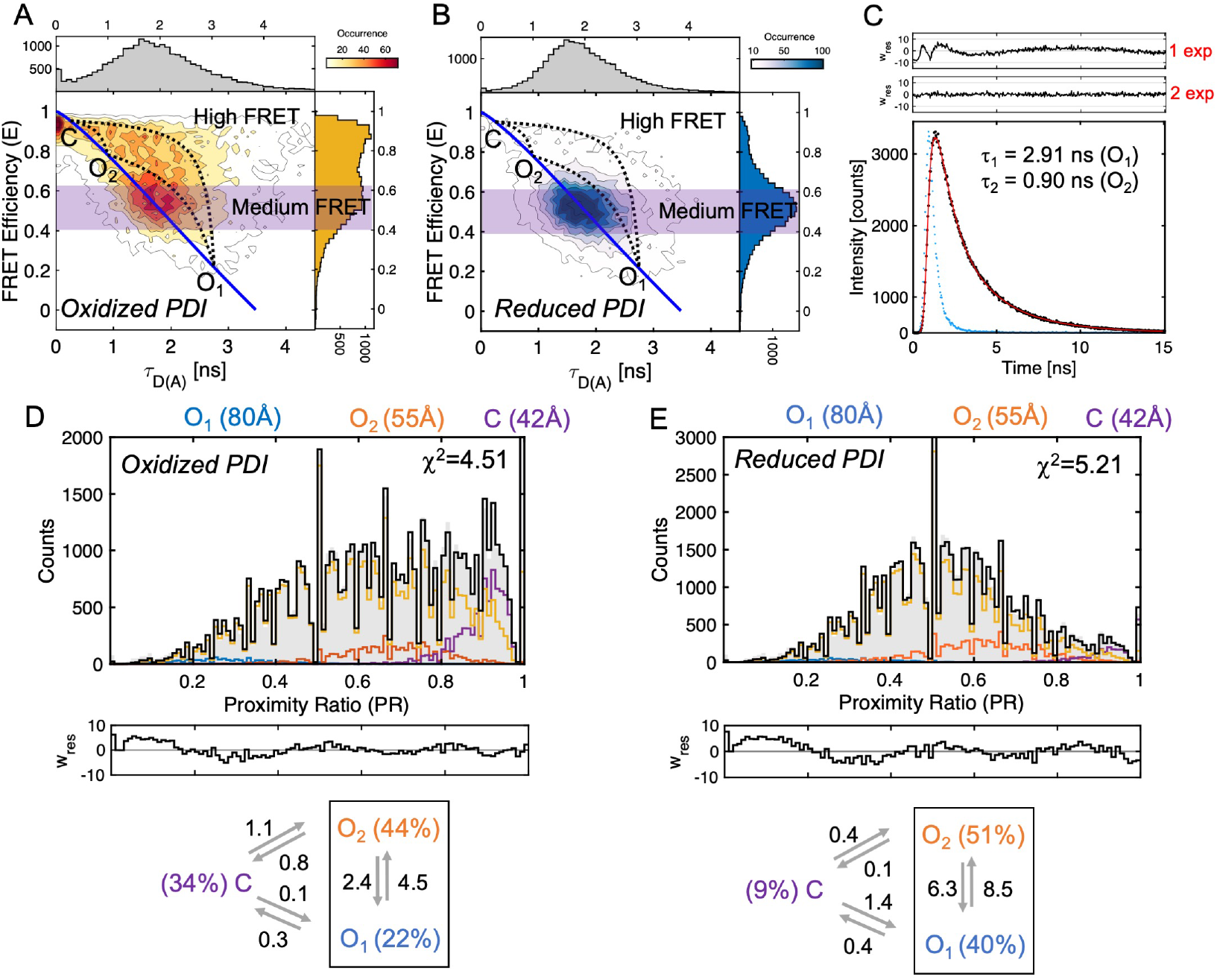
Dynamics of oxidized and reduced PDI. 2D plots of FRET efficiency versus lifetime of the donor in the presence of the acceptor τ_D(A)_ for **(A)** oxidized and **(B)** reduced PDI 88/467 documenting the dynamic exchange between closed (high FRET) and open (medium FRET) ensembles. The solid blue line describes the theoretic relationship between FRET and lifetime (“static FRET line”). Systematic deviations from this relationship highlighted by the dotted lines track the trajectory of single molecules that are exchanging between C, O_1_ and O_2_ while passing through the confocal volume. The magenta regions indicate the molecules selected for the lifetime analysis (0.4-0.6) shown in **panel C. C)** Subpopulation specific fluorescence lifetime of oxidized PDI 88/467. Data points are in black. The red line represents the best fit obtained with a double exponential function (χ^2^=1.33). The value of lifetime of each population is shown in the plot. It is also reported in **Table 3** together with the amplitude for each population. The instrumental response function is shown in blue. Weighted residuals for one (1 exp) and two exponential (2 exp) fit are shown above the graph. PDA analyses of oxidized **(D)** and reduced **(E)** PDI 88/467 obtained with a dynamic three-state model. C, O_1_ and O_2_ sates are shown in purple, orange and blue, respectively while the exchange between them is represented by the yellow line. The black line represents the global fit. Weighted residuals are shown below each plot together with a diagram that summarizes the kinetic scheme, the rate constants and fraction of each population at equilibrium. Note how the residuals show a larger deviation toward lower values of PR. Such deviation is most likely due to acceptor fluorophore that goes into a dark, non-FRET state.

In contrast to the closed population, the center of the open population did not reside on the static FRET line but was instead slightly shifted toward the right. This was particularly evident for oxidized PDI 88/467. Since the dyes are freely rotating in solution thus not theoretically affecting protein dynamics, we hypothesized that, in addition to C, PDI visits multiple FRET states within the open ensemble that exchange in the millisecond timescale. This is because PDI molecules remain, on average, ∼0.5 ms in the observation volume.

**Table 3.** Subpopulation specific (0.4-0.6) fluorescence lifetime analysis of oxidized and reduced PDI 88/467.

| | $\tau_1$ (ns) | f1 | $\tau_2$ (ns) | f2 | $\chi^2$ |
| --- | --- | --- | --- | --- | --- |
| **oxidized 88-467** |  |  |  |  |  |
| **1 exp** | $2.23 \pm 0.08$ | 1 | - | - | 9.69 |
| **2 exp** | $2.91 \pm 0.15$ | 0.44 | $0.90 \pm 0.06$ | 0.56 | 1.32 |
| **reduced 88-467** |  |  |  |  |  |
| **1 exp** | $2.04 \pm 0.08$ | 1 | - | - | 14.58 |
| **2 exp** | $2.72 \pm 0.15$ | 0.42 | $0.91 \pm 0.06$ | 0.58 | 1.32 |

Values of lifetime are indicated by $\tau$ . The fraction of each population (0-1) is indicated by f. The quality of the fit is expressed by the value of $\chi^2$ . The same dataset was fit with one, two and three exponential functions using the lifetime fit module in PAM. Satisfactory fit was attained for both oxidized and reduced PDI with a two exponential decay model. Since no further improvement was observed with a three exponential models, this result was discarded and not presented in the Table.

To test this hypothesis, we performed subpopulation specific fluorescence lifetime analysis, a methodology that has proven useful to define the number of species at equilibrium that are faster than diffusion *(23, 25, 31)*. Since we hypothesized heterogeneity within the open ensemble, we selected bursts from the FRET interval 0.4-0.6, which is highlighted in magenta. If multiple PDI species were present at equilibrium, we expected more than one relaxation would be necessary to fit the lifetime plots. For both oxidized and reduced PDI 88/467 **(Figure 3C** and **Table 3)**, the lifetime decay could not be fit with one exponential (1 exp, χ^2^=9.69) but instead required a double exponential function (2 exp, χ^2^=1.31). The addition of a third relaxation did not significantly improve the fit (χ^2^=1.28). This result agrees with our hypothesis and documents the existence of two states within the open ensemble, which we called O_1_ and O_2_. Additionally, we found that the ratio f_2_/f_1_ between the amplitudes f_2_ for τ_2_ and f_1_ for τ_1_ was ∼1.2 in both oxidized and reduced PDI. We concluded that O_2_ is more represented at equilibrium, regardless of the redox state.

Previous studies have suggested that oxidized and reduced PDI are structurally distinct *(16, 20, 32)*. Despite the macroscopic differences between the FRET profiles, we, however, measured values of τ_1_ and τ_2_ that are very similar, within experimental error **(Table 3)**. This indicates that very similar, perhaps identical, conformational states exist in both oxidized and reduced PDI. Given PDI’s flexibility, we propose that the redox microenvironment modulates PDI by a conformational selection mechanism rather than forming new macroscopic species.

**Connectivity between the ensembles and rate of interconversion**

By having identified the minimum number of macroscopic states at equilibrium, we next defined the connectivity between them *(23, 29)*. To this end, using the previously determined values of lifetime of 0.25 ns for C, 0.91 ns for O_2_ and 2.75 ns for O_1_, which, using the equation E=1-(τ_D(A)_/τ_D_), a τ_D_=3.5 ns and an R_0_=65Å, correspond to E_1_=0.93 (or 42Å) for C, E_2_=0.73 (or 55Å) for O_2_ and E_3_=0.22 (or 80Å) for O_1_, we drew the corresponding dynamic FRET lines (dotted lines) in **Figure 3A** and **Figure 3B**. These lines were drawn according to previous work in the field *(23–25, 30, 33)*. Although less evident for reduced PDI because of the low intensity of C, we found bursts lying on all three lines indicating dynamic exchange between the FRET states. This led us to propose a triangular kinetic scheme for both reduced and oxidized PDI, implying that, in solution, the three ensembles spontaneously exchange to one another.

To quantify the abundance of each ensemble at equilibrium and determine the rates at which they interconvert, we performed photon distribution analysis (PDA) *(25, 33, 34)*. Given the results of our previous experiments, we chose a three-state dynamic model. Datasets for oxidized and reduced PDI 88/467 binned at 1, 0.75, 0.5 and 0.25 ms were globally fit after fixing the value for each state to 42Å, 55Å and 80Å, which are the experimentally determined values for C, O_2_ and O_1_ **(Figure S6)** and restricting the value of sigma (σ) to 0.045. Sigma defines the width of a shot-noise limited distribution and was experimentally determined in our system using fluorescently labeled double stranded DNA constructs with different lengths **(Figure S7)**. The value of 0.045 also agrees with the results of recent studies aimed at comparing the accuracy and reproducibility of smFRET data among multiple laboratories *(35)*. Representative results obtained with datasets binned at 0.5 ms are shown in **Figure 3D** and **Figure 3E** for oxidized and reduced PDI 88/467, respectively. The rate constants measured by PDA are reported in **Table 4** and summarized in the scheme located below each plot.

**Table 4.**
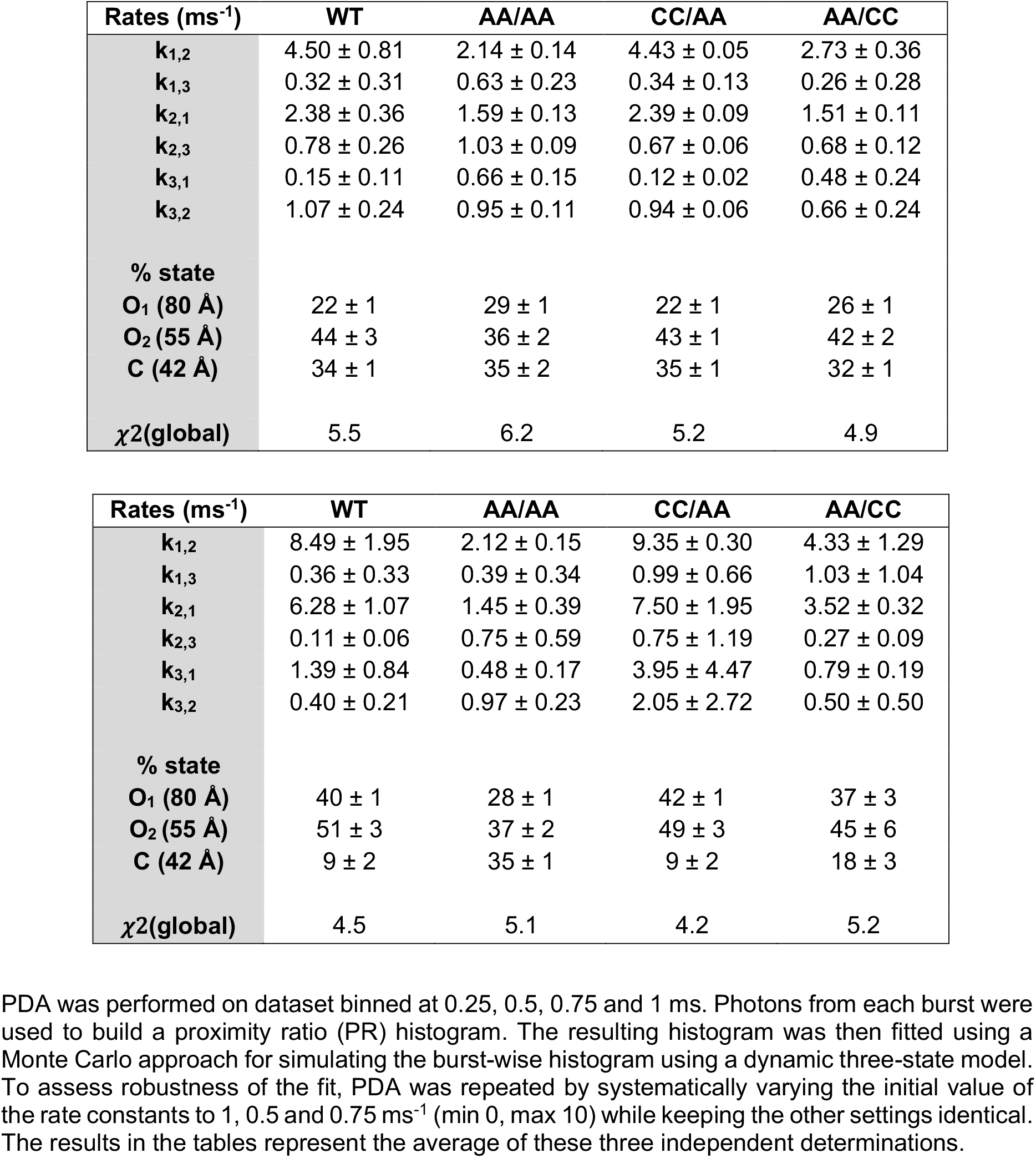
PDA analysis of oxidized (top) and reduced (bottom) PDI.

| **Rates (ms<sup>-1</sup>)** | **WT** | **AA/AA** | **CC/AA** | **AA/CC** |
| --- | --- | --- | --- | --- |
| **k<sub>1,2</sub>** | 4.50 ± 0.81 | 2.14 ± 0.14 | 4.43 ± 0.05 | 2.73 ± 0.36 |
| **k<sub>1,3</sub>** | 0.32 ± 0.31 | 0.63 ± 0.23 | 0.34 ± 0.13 | 0.26 ± 0.28 |
| **k<sub>2,1</sub>** | 2.38 ± 0.36 | 1.59 ± 0.13 | 2.39 ± 0.09 | 1.51 ± 0.11 |
| **k<sub>2,3</sub>** | 0.78 ± 0.26 | 1.03 ± 0.09 | 0.67 ± 0.06 | 0.68 ± 0.12 |
| **k<sub>3,1</sub>** | 0.15 ± 0.11 | 0.66 ± 0.15 | 0.12 ± 0.02 | 0.48 ± 0.24 |
| **k<sub>3,2</sub>** | 1.07 ± 0.24 | 0.95 ± 0.11 | 0.94 ± 0.06 | 0.66 ± 0.24 |
| **% state** |  |  |  |  |
| **O<sub>1</sub> (80 Å)** | 22 ± 1 | 29 ± 1 | 22 ± 1 | 26 ± 1 |
| **O<sub>2</sub> (55 Å)** | 44 ± 3 | 36 ± 2 | 43 ± 1 | 42 ± 2 |
| **C (42 Å)** | 34 ± 1 | 35 ± 2 | 35 ± 1 | 32 ± 1 |
| **χ<sup>2</sup>(global)** | 5.5 | 6.2 | 5.2 | 4.9 |
| **k<sub>1,2</sub>** | 8.49 ± 1.95 | 2.12 ± 0.15 | 9.35 ± 0.30 | 4.33 ± 1.29 |
| **k<sub>1,3</sub>** | 0.36 ± 0.33 | 0.39 ± 0.34 | 0.99 ± 0.66 | 1.03 ± 1.04 |
| **k<sub>2,1</sub>** | 6.28 ± 1.07 | 1.45 ± 0.39 | 7.50 ± 1.95 | 3.52 ± 0.32 |
| **k<sub>2,3</sub>** | 0.11 ± 0.06 | 0.75 ± 0.59 | 0.75 ± 1.19 | 0.27 ± 0.09 |
| **k<sub>3,1</sub>** | 1.39 ± 0.84 | 0.48 ± 0.17 | 3.95 ± 4.47 | 0.79 ± 0.19 |
| **k<sub>3,2</sub>** | 0.40 ± 0.21 | 0.97 ± 0.23 | 2.05 ± 2.72 | 0.50 ± 0.50 |
| **% state** |  |  |  |  |
| **O<sub>1</sub> (80 Å)** | 40 ± 1 | 28 ± 1 | 42 ± 1 | 37 ± 3 |
| **O<sub>2</sub> (55 Å)** | 51 ± 3 | 37 ± 2 | 49 ± 3 | 45 ± 6 |
| **C (42 Å)** | 9 ± 2 | 35 ± 1 | 9 ± 2 | 18 ± 3 |
| **χ<sup>2</sup>(global)** | 4.5 | 5.1 | 4.2 | 5.2 |

The most notable difference between the two redox sates identified by PDA concerns the distribution of the three ensembles at equilibrium. In the presence of DTT **(Figure 3E)**, C was minimally populated (∼9%) whereas O_1_ and O_2_ accounted for ∼40% and ∼51%, respectively. By contrast, oxidized PDI spent similar amount of time in C and O_1_ but preferred O_2._ The fact that O_2_ dominates agrees with our previous analysis and suggests that O_2_ is the preferred state adopted by unbound PDI in solution, regardless of the redox state.

Other differences between the two redox sates concerned the magnitude of the rate constants. We found that the rate at which O_2_ converts to C (k_2,3_) was ∼8 time faster for oxidized PDI compared to reduced PDI. In contrast, the rate at which C converts to O_1_ (k_3,1_) was ∼9 times slower in oxidized PDI compared to reduced PDI. Faster O_2_→C conversion and slower C→O_1_ conversion explain why more C is present in oxidized PDI compared to reduced PDI. We also found that the transition O_1_⟷O_2_ was the fastest of the catalytic cycle, and significantly faster (∼6 fold) than diffusion, especially for reduced PDI. This latter observation is important for two reasons. First, it explains why O_1_ and O_2_ cannot be individually visualized in the 2D plots of FRET efficiency versus lifetime but instead merge to form a broad ensemble. Second, it predicts that, when PDI dwells in either O_1_ or O_2_, transition to C is energetically more expensive, especially when PDI is reduced. From a structural standpoint, this indicates that O_1_ and O_2_ may be alike, yet significantly different from C.

To further confirm that O_1_ and O_2_ exchange rapidly, we performed species selected filtered fluorescence correlation spectroscopy (fFCS). In this method, as described by Felekyan et al. *(36)*, auto-correlation and cross-correlation functions are calculated for two species of interest to determine the presence of dynamic exchange between them and the interconversion rates. fFCS analysis was performed between O_1_ (0.17<E<0.25) and O_2_ (0.65<E<0.75) in oxidized and reduced states. The results are shown in **Figure 4A** and **Figure 4B**.

**Figure 4.**
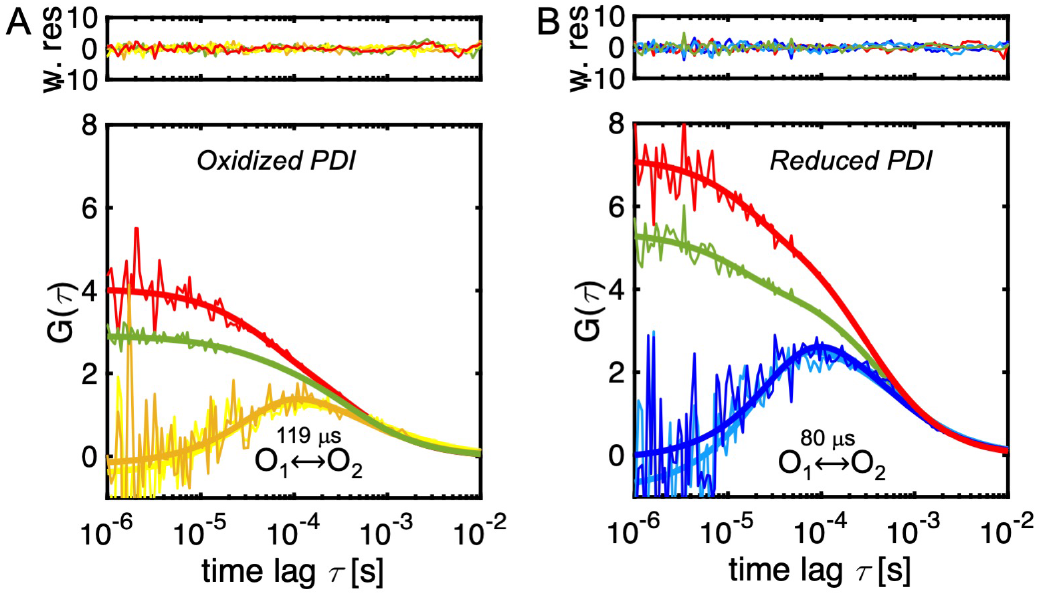
Rapid exchange between O_1_ and O_2_ ensembles monitored by fFCS. Auto-correlation (red and green) and cross-correlation curves (yellow and blue) of O_1_ and O_2_ ensembles for oxidized **(A)** and reduced **(B)** PDI 88/467. The solid lines represent the best fit model obtained by globally fitting the four curves, which enables extraction of the interconversion time, expressed in microseconds. Randomly distributed weighted residuals are shown above each plot. Best fit parameters are τ_R,ox =_ 119±12 μs; τ_D =_ 469 μs (fixed, diffusion), χ^2^=1.11; τ_R,rd =_ 80±9 μs; τ_D =_ 469 μs (fixed, diffusion), χ^2^=1.25.

The presence of a bell-shaped cross-correlation function between O_1_ and O_2_ documents rapid exchange occurring on a time scale comparable or faster than the diffusion time. Global fitting of the four correlation curves (two sACFs and two sCCFs) required, in addition to the diffusion term (τ_D_), an additional relaxation term, τ_R_, providing conclusive evidence of fast dynamics. After fixing the diffusion term to 469 μs (see methods), the values of τ_R_ calculated for oxidized and reduced PDI 88/467 were 119±12 μs and 80±9 μs, respectively. These values are in reasonable agreement with the rate constants measured by PDA for the O_1_⟷O_2_ exchange (τ_R_=(k_1,2_ + k_2,1_)^-1^), which are 145 μs for oxidized PDI and 67 μs for reduced PDI, respectively. Thus, fFCS successfully visualized rapid exchange between O_1_ and O_2_ and confirmed that one of the main differences between oxidized and reduced PDI is the ability of O_1_ and O_2_ to exchange faster in reduced PDI but slower in oxidized PDI.

### Nonequivalent structural role of the active sites

PDI has two active sites, one in the N-terminal **a** domain (C53/C56) and another one in the C-terminal **a’** domain (C397/C400). To address how the four catalytic cysteines control the newly discovered conformational equilibrium, they were mutated to the redox-insensitive amino acid alanine (A) to generate the three new constructs, namely the quadruple mutant PDI 88/467/C53A/C56A/C397A/C400A (PDI-AA/AA), and the double mutants PDI 88/467/C53A/C56A (PDI-AA/CC) and PDI 88/467/C397A/C400A (PDI-CC/AA). After verifying that the mutants had activity profiles consistent with what was previously reported in the literature for the wild-type background *(37)* **(Figure S8)**, smFRET measurements were collected in the absence and presence of DTT. PDA analysis was used to quantify the species at equilibrium as described previously. Results of PDA analysis are reported in **Table 4**. To facilitate comparison between PDI 88/467 wild-type and mutants, 1D FRET efficiency plots are shown the main text **(Figure 5)**. 2D FRET efficiency vs. lifetime plots are shown in the supplementary materials **(Figure S9)**.

**Figure 5.**
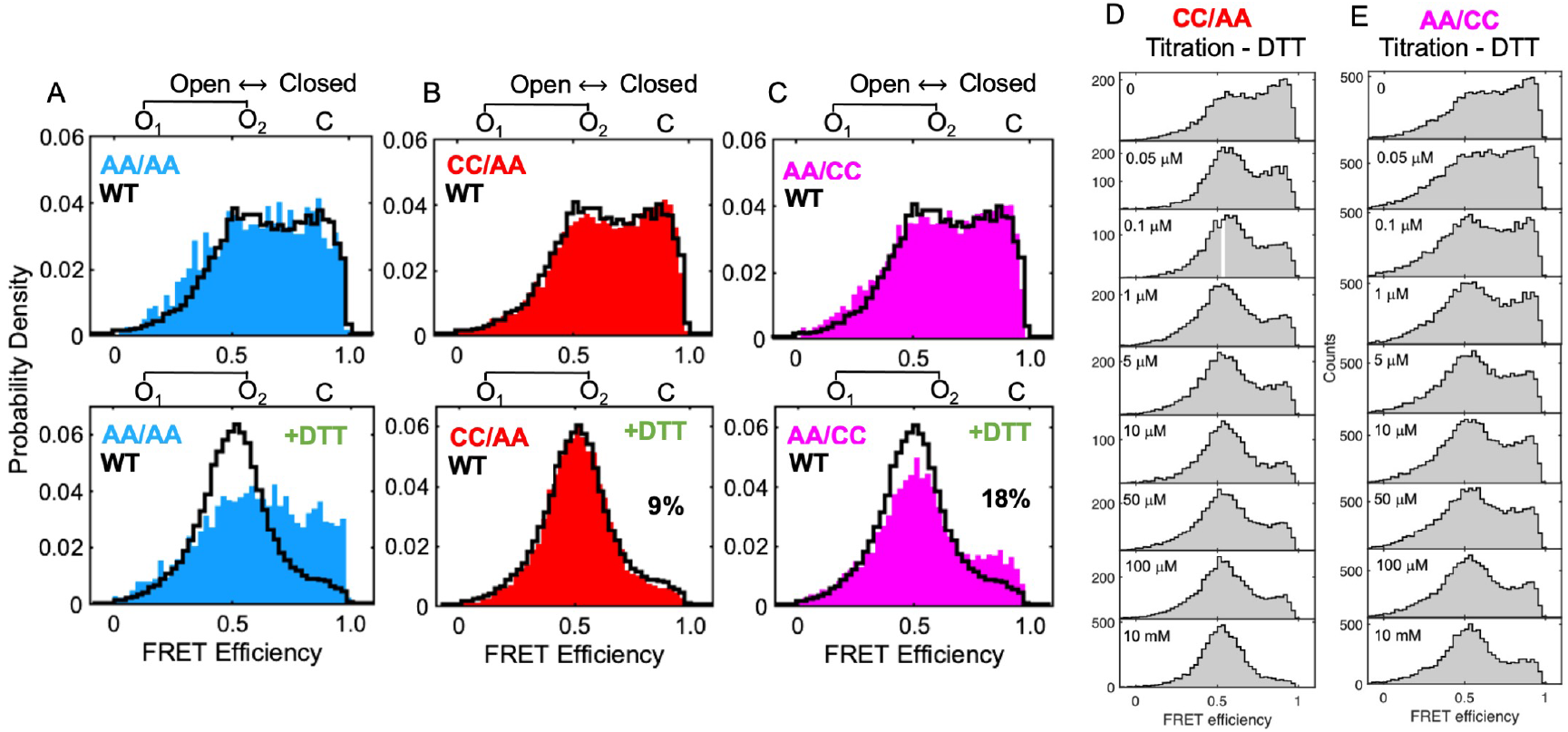
Structural nonequivalence of the active sites. Normalized 1D FRET efficiency histograms of **(A)** PDI 88/467 C53A/C56A/C397A/C400A (AA/AA, blue), **(B)** PDI 88/467 C397A/C400A (CC/AA, red) and **(C)** PDI C53A/C56A (AA/CC, magenta) overlaid to PDI WT (black) before (top panel) and after (bottom panel) the addition of 1 mM DTT. Note how PDI AA/AA is similar to oxidized PDI 88/467 wild-type (WT) and insensitive to DTT and how the mutations C53A and C56A led to a macroscopic accumulation of C, which, according to PDA analysis **(Table 4)** increased 2-times, from 9% to 18%. 1D FRET efficiency histograms of PDI 88/467 C397A/C400A (CC/AA) **(D)** and PDI C53A/C56A (AA/CC) **(E)** collected at increasing concentrations of DTT (0-10 mM), covering 4 orders of magnitudes.

A first important finding was that, in the absence of DTT, the active site variants were similar to each other and also similar, yet not identical, to oxidized PDI 88/467 **(Figure 5, top row)**. This data indicates that the catalytic cysteines are not required for initiating large-scale domain movements such as those monitored here by smFRET. These domain motions must, therefore, occur spontaneously, favored by the flexibility of the protein fold. It is important to point out, however, that, even though the equilibrium distribution of C, O_1_ and O_2_ remained mostly unchanged, the rates at which the three states exchanged slightly decrease compared to PDI 88/467 **(Table 4)**, indicating that the catalytic cysteines are important for protein dynamics.

Another important observation was that the active site mutants, while similar in the oxidized state, behaved differently in the presence of DTT **(Figure 5, bottom row)**. Specifically, PDI-AA/AA was insensitive to the addition of DTT; PDI-CC/AA behaved just like PDI 88/467; and PDI-AA/CC was in between PDI-AA/AA and PDI-CC/AA insofar as it partly responded to the addition of DTT, which led to a significant twofold accumulation of C **(Figure 5C, bottom row)**. Importantly, this effect was not due to a reduced reactivity of the mutant toward DTT, rather to changes in protein dynamics caused by the mutations. This is because C never disappeared, even at very high (10 mM) concentrations of DTT **(Figures 5D and 5E)**. We concluded that: 1) the active site cysteines are responsible for sensing the redox microenvironment and 2) the N- and C-terminal active sites are nonequivalent in the context of PDI dynamics.

### Active site ligation stabilizes closed conformations of PDI

To further investigate the role of the active sites in controlling the allosteric equilibrium, we took advantage of their reactivity toward the commercially available inhibitor 16F16 *(38)*, which contains a chloroacetamide electrophile for covalent modification of PDI. In contrast to commonly used alkylating agents such as N-ethylmaleimide (NEM) and diamide, 16F16 is more potent and specifically react with the active site cysteines C53 and C397 *(39)*. Furthermore, at the concentration used in this study, 16F16 does not quench the fluorescence intensity of the Atto dyes, which was significantly compromised with mM concentrations of NEM and diamide. The results of this experiment are shown in **Figure 6A** and **Figure 6B**. Starting from a solution of oxidized PDI 88/467, we added, in a sequential order, 50 µM DTT and then, after 70 min, 50 μM 16F16. Addition of DTT and 16F16 are indicated with red arrows. This design enabled us to follow in real-time PDI conformational cycle. We found that addition of 16F16 to reduced PDI shifted the equilibrium toward the closed ensemble in a time-dependent fashion. Since 16F16 did not change the FRET profile of oxidized PDI *(data not shown)*, we interpreted this as evidence that binding of 16F16 followed by alkylation of the catalytic cysteines stabilizes closed conformations of PDI. Similar results were obtained with another commonly used irreversible active site inhibitor of PDI, namely PACMA-31 **(Figure 6C)** *(40)*, arguing for a general mechanism of this class of compounds.

**Figure 6.**
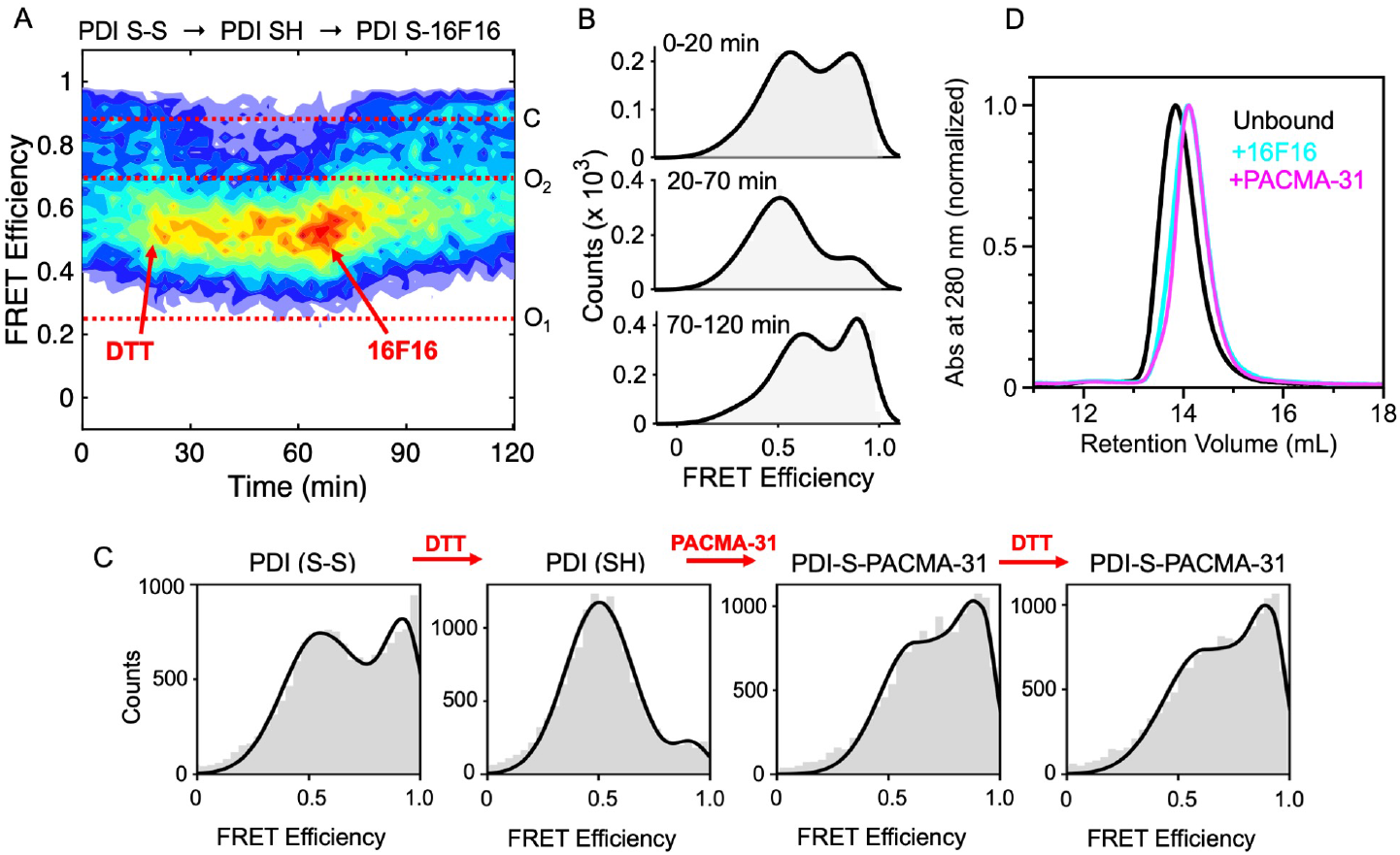
Active site ligation stabilizes closed conformations of PDI. A solution of PDI 88/467 (50 pM) was continuously monitored for 120 min under different experimental conditions. Addition of 50 μM DTT and 50 μM 16F16 is indicated with red arrows. The horizontal dotted lines identify the mean FRET efficiency value of C, O_1_ and O_2_. **B)** FRET efficiency histograms of PDI 88/467 at three different time intervals monitoring key steps of the reaction of PDI 88/467 with 16F16. **C)** FRET histograms of PDI 88/467 (100 pM, TBSE-T) in the absence and presence of DTT (1 mM) before and after the addition of 50 μM PACMA-31. Note how PACMA-31, like 16F16, shifts the conformational equilibrium towards the closed ensemble. Further addition of DTT is inconsequential. This is because PACMA-31 reacts irreversibly with the active site thiol groups of PDI to form a covalent adduct. **D)** SEC analysis of PDI free and bound to 16F16 (cyan) and PACMA-31 (magenta).

To independently validate this observation, we performed SEC experiments using PDI wild-type free and bound to 16F16 and PACMA-31 **(Figure 6D)**. The retention volume of PDI was delayed of ∼0.4 ml in the presence of 16F16 and PACMA-31 compared to unbound PDI. Since proteins with smaller hydrodynamic radius have larger retention volumes, this result supports the compaction model upon ligation inferred by smFRET data.

## Discussion

This study reports a detailed biophysical characterization of PDI’s dynamics in solution and identifies several new features of this allosteric enzyme that were not known before. Using a combination of four novel FRET pairs located in the **a** and **a’** catalytic domains and three active site mutants, we discovered that PDI visits, on the sub-millisecond timescale, three major conformational ensembles at equilibrium, O_1_, O_2_ and C, whose distribution is regulated by a variety of factors, namely the redox microenvironment **(Figure 2)**, the presence of active site cysteines **(Figure 5)** and active site ligands **(Figure 6)**. Importantly, the identification of these ensembles is fully consistent with previous structural, biophysical, and biochemical data documenting structural flexibility of the protein fold *(16, 18–20, 32)*. It thus represents an important step forward for achieving a deeper mechanistic understanding of how this enzyme works under physiologically relevant conditions.

While PDI’s flexibility was expected, the discovery that the microenvironment modulates PDI by a conformational selection mechanism is conceptually new. As such, this finding enables us to propose a new model of PDI dynamics whereby environmental factors commonly found in cells and in the circulation, such as high levels of levels of oxidative stress and chemical modifications of catalytic cysteines (i.e., sulfenylation, S-nitrosylation and acylation) affect the solution structure of PDI by shifting this equilibrium without forming new macroscopic species. Selection between pre-existing PDI ensembles provides the structural basis for understanding how PDI activity is regulated by conformational modulation.

Also new and illuminating is the finding that, despite undergoing large conformational changes, O_1_, O_2_ and C, interconvert very rapidly. Energetically, this indicates that the free-energy landscape of PDI is characterized by low basins and shallow minima. To visualize this, we calculated free-energy barriers for transition between the ensembles using the Arrhenius equation and the reaction rate constants reported in **Table 4 (Figure 7)**. Quite remarkably, these calculations yielded Gibbs free-energy values (ΔG) lower than 10K_B_T, which is the energy required to form/break less than two hydrogen bonds (6.7K_B_T is the energy calculate for one hydrogen bond using the same pre-exponential factor used by us and others *(41)*). Considering the magnitude of the conformational changes monitored by smFRET and the low free-energy barriers required for transitioning between the states, we propose that, in solution, PDI’s flexibility arises from rapid relocations of the domains mediated by the linkers. To satisfy these energetic requirements, the linkers, however, must be free to move, thus only weakly interacting with the surrounding domains. While more studies are needed to validate this model, recent results obtained in our laboratory with the FRET pair 308/467 *(22)*, in which dyes are located across linker 3, also known as the x-linker, supports this view. In fact, PDI 308/467 not only displays multiple FRET states at equilibrium but also displays kinetic features that are similar, yet not identical, to the one reported here for FRET pairs located in the **a** and **a’** domains. Considering these new findings, alternative models predicting the x-linker to mediate PDI dynamics by binding and dissociating with a hydrophobic pocket in **b′** *(42-44)* should be reconsidered.

**Figure 7.**
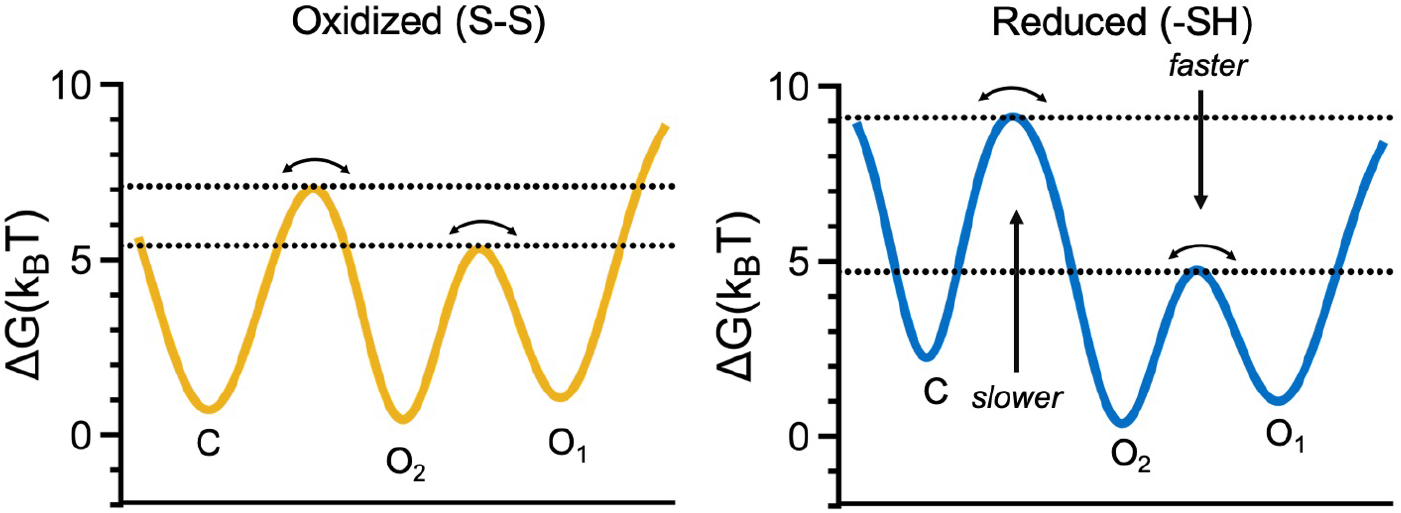
Conformational landscape of PDI inferred by smFRET. Free energy profiles of oxidized (left, yellow) and reduced (right, blue) PDI obtained using data reported in **Table 4**. Barrier heights corresponding to C→O_2_ and O_1_→O_2_ (horizontal lines) for oxidized and reduced PDI were calculated using the Arrhenius equation with a pre-exponential factor of 10^5^ s^−1^. k_B_ is the Boltzmann constant and T is the temperature. The distributions were arbitrarily drawn using a combination of Gaussian distributions. Note how in the presence of DTT the free-energy barrier increases for the C⟷O_1_/O_2_ transition (slower transition) whereas it decreases for the O_1_⟷O_2_ transition (faster transition) leading to redistribution of the conformational ensembles.

Another interesting observation emerging form our smFRET analysis of PDI was that the FRET profile of the redox-insensitive variant PDI AA/AA was similar to the FRET profile of oxidized PDI, not reduced PDI. This result was unexpected based on previous literature *(16, 20)* and indicates that, while active site thiols are necessary to drive conformational changes in PDI, the presence of disulfide bonds is not. While the structural basis behind this observation cannot be inferred by these studies, because of the different pKa values between the two cystine residues of the catalytic motif *(1)*, we speculate that protonation of the resolving cysteine may be key to initiate this process.

Finally, PDI interacts with many substrates intracellularly and extracellularly. The mechanisms of substrate recognition and release are not fully understood. To date, the most popular model for PDI-assisted catalysis is based on X-ray crystal data *(16)* and envisions oxidized PDI adopting a flexible open form, which is primed for substrate binding. After transferring the disulfide bond to the substrate, reduced PDI is then believed to become more compact and rigid, thus favoring substrate release *(16, 20, 32)*. Evaluation of PDI by smFRET led to identification of new structural features that challenge this model. In contrast with what was predicted by X-ray crystallography, our data indicate that residues 57/467 and residues 88/467 spend significantly more time away from each other in reduced PDI compared to oxidized PDI. In solution, the catalytic domains of reduced PDI may therefore visit conformations that are significantly more open than what has been captured by X-ray crystallography. At the same time, residues 57/467 and residues 88/467 spend significantly more time close to each other in oxidized PDI compared to reduced PDI. On these bases, we propose an alternative conformational cycle for PDI whereby oxidized PDI can adopt very compact conformations upon substrate binding and, therefore, substrate release is facilitated by opening, not closing, of the structure. This mechanism agrees well with findings in **Figure 6** documenting that active site ligation favors closed conformations of PDI. It could also explain how reduced PDI, which is, on average, more open than oxidized PDI, can easily interact and process with very bulky substrates such as clotting factors.

## Materials and Methods

### Protein production and purification

The cDNA of human PDI (residues 18-479) was cloned into a pBAD vector expression system (ThermoFisher) and modified to include an N-terminal 6 his-tag and a C-terminal Avitag. Genetic incorporation of the unnatural amino acidic N-Propargyl-L-Lysine (Prk) (SiChem) at positions 57, 88, 401 and 467 was obtained using the AMBER suppressor pyrrolysine tRNA/RS system from *Methanosarcina mazei*. Mutations C53A, C56A, C397A, C400A in the 88/467 background were generated using the Quickchange Lightning kit (Agilent) with appropriate primers. Sequence verified PDI variants (Genewiz) were expressed in Top10 cells and purified following recently published procedures *(22)*.

### Protein labeling

Site-specific labeling was achieved as detailed elsewhere *(22)*. Briefly, a solution of 25 μM of PDI in 1x phosphate buffer saline (PBS) pH 7.4 (Corning) was reacted with 4x molar excess of azide dyes (donor, acceptor or donor/acceptor mixtures) (Sigma-Aldrich) in the presence of 150 μM copper sulfate (CuSO_4_), 750 μM tris-hydroxypropyltriazolylmethylamine (THPTA), and 5 mM sodium ascorbate. The reaction mixer was left on slow rotisserie for 1 hour 30 minutes at room temperature, then 30 minutes on ice. The reactions were stopped by adding 5mM EDTA. Monomeric PDI was successfully separated by protein aggregates by size exclusion chromatography (SEC), using a Superdex 200 10/300 column (Cytiva) equilibrated with Tris 20 mM (pH 7.4), 150 mM NaCl, 2 mM EDTA. The quality of each protein preparation was assessed by NuPAGE Novex 4–12% Bis-Tris protein gels (ThermoFisher). Gels were stained with Coomassie Brilliant Blue R-250 (ThermoFisher) and scanned on a Typhoon imager (Cytiva) at 532 nm and 633 nm to verify specific incorporation of the fluorescent dyes. Total protein concentration was determined by reading the absorbance at 280, using a molar coefficient adjusted for the amino acidic sequence of each variant. The concentration of Atto-550 and Atto-647N was calculated by reading the absorbance at 550 nm and 640 nm, respectively. Typical labeling efficiencies were 90%.

### Circular Dichroism (CD)

Far-UV CD spectra were recorded on Jasco J-715 spectropolarimeter equipped with a water-jacketed cell holder, connected to a water-circulating bath, as done before *(22)*. Spectra were collected for unlabeled and labeled protein in PBS with 2mM EDTA at a concentration of 0.12 mg/ml. The final spectra resulted from the average of five accumulations after base line subtraction.

### Intrinsic Fluorescence Assay

Intrinsic fluorescence spectra (tryptophan) were performed in a reaction volume of 200 μl with 0.2 μM of PDI in 20 mM Tris-HCl buffer containing 150 mM NaCl (pH 7.4) and either 1mM GSH or 1mM GSSG were incubated for 1hr at room temperature. Emission spectra were recorded at 295–450 nm with excitation at 280 nm using a FluoroMax-4 (Horiba).

### Insulin reductase assay

PDIs (400 nM) were solubilized in PBS and then added to a solution containing 0.2 mM human insulin (Sigma-Aldrich), 2 mM EDTA and 325 μM DTT. The reaction was monitored at 650 nm (turbidity due to precipitation of the product) for 1 h at 25 °C using a Spectramax i3 (Molecular Devices). Statistical analysis was performed using unpaired t-test in Prims 9.0.

### Determination of Anisotropy and Quantum Yield

Four singly labeled PDI constructs (i.e., K57U, S88U, K401U and K467U) were expressed, labeled with either Atto-550 or Atto-647N and purified as described before for anisotropy and quantum yield determination. Anisotropy was recorded in 1x PBS with 2 mM EDTA buffer using Fluorolog-3 (Jobin-Yvon). The Atto-550 labeled proteins (50 nM) were excited at 540 nm and emission was monitored at 580 nm. The Atto-647N labeled proteins (50nM) were excited at 640 nm and emission was monitored at 680 nm, excitation and emission slits were set at 1 and 14 nm, respectively. The donor quantum yield was measured in bulk fluorescence assays in a FluoroMax-4 (Jobin–Yvon) for each donor position, in reference to the quantum yield of Rhodamine 110 (89.87+/-0.91). The emission spectra of PDI labeled with only a donor at positions 57, 88, 401 and 467 were collected at five concentrations under the same excitation conditions (532 nm) in the buffer used for smFRET experiments. The quantum yield was found from the ratio between the dye’s integrated emission spectrum and its absorbance at 532 nm. The overlap between the donor emission spectrum and the acceptor absorbance spectrum is defined as:

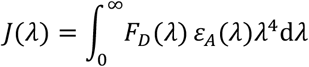

Where F_D_ (λ) is the normalized donor emission spectrum, and ε_A_ is the acceptor’s absorbance spectrum, measured for PDI labeled with an acceptor at position at position 57, 88, 401 and 467 respectively.

### Single-molecule FRET measurements

FRET measurements of freely diffusing single molecules were performed with a confocal microscope MicroTime 200 (PicoQuant) using published procedures *(22, 45, 46)*. Excitation laser light from 532 nm and 638 nm lasers was used to excite the donor and acceptor fluorophores, respectively. A Pulsed Interleaved Excitation (PIE) setup was used with a pulse rate of 20 MHz to alternate the donor and acceptor excitation. PIE reports the status of both donor and acceptor fluorophores by sorting molecules based on relative donor:acceptor stoichiometry (S) and apparent FRET efficiency (E), as described before *(24, 30, 47, 48)*. A dichroic mirror (ZT405/488/532/640rpc-XT, Chroma) reflecting at 532 and 638 nm guided the light to a high numerical aperture apochromatic objective (60x, N.A. 1.2, water immersion, Olympus) that focused the light to a confocal volume of 1.0 fl. Fluorescence from excited molecules was collected with the same objective and focused onto a 50-μm diameter pinhole. The donor and acceptor emissions were separated via a dichroic filter with a dividing edge at 620 nm (620DCXR, Chroma). Suited bandpass filters (HQ580/70m, Chroma and HQ690/70m, Chroma) were inserted to eliminate the respective excitation wavelength and minimize spectral crosstalk. The fluorescence was detected with two single-photon avalanche diode detectors (τ-SPAD, Perkin Elmer) using Time-correlated Single Photon Counting with the TimeHarp 200 board (HydraHarp 400, PicoQuant). Data were stored in the Time-tagged Time-resolved Mode as a PTU file format.

Measurements were performed 25 μm deep in the solution using a laser power of ∼15 μW. Total acquisition time was ∼40 minutes per sample. Data collection was repeated for a minimum of four times using the same sample as well as new protein samples from at least one different preparation. Identical results were obtained in all cases, indicating stability of the protein sample during data collection and reproducibility. Concentration was 50-100 pM of labeled protein solubilized in 200 μl of 20 mM Tris, 150 mM NaCl, 2 mM EDTA, 0.003% Tween 20, pH 7.4 (TBSE-T) for oxidized PDI and TBSE-T with 1 mM DTT for reduced PDI. DTT titrations were performed by preincubating oxidized PDI with the desired concentration of DTT for 40 minutes at room temperature before data collection to ensure equilibrium. However, similar results were obtained by adding increasing concentrations of DTT to the same protein sample, implying fast reactivity of DTT towards the enzyme. The PDI inhibitors 16F16 (Sigma-Aldrich) and PACMA-31 (Sigma-Aldrich) solubilized in DMSO were added to the solution at a final concentration of 50 μM. The inhibitory effect of 16F16 and PACMA-31 was verified in the insulin reductase assay, as detailed above. Data recording was performed using the Sympho-Time Software 64, version 2.3 (PicoQuant, Berlin).

### Single-molecule FRET analysis

Data analysis was carried out with the Matlab-based software PAM *(49)* using a customized profile optimized for our microscope. Signals from single molecules were observed as bursts of fluorescence. Bursts with more than 40 counts were searched with the All Photon Burst Search (APBS) algorithm. Integration time was set to 0.5 ms. Appropriate corrections for direct excitation of the acceptor at the donor excitation wavelength (DE), leakage of the donor in the acceptor channel (Lk), and the instrumental factor (γ) were determined experimentally using a mixture of double-stranded DNA models with known FRET efficiency and stoichiometry labeled with dyes Atto-550 and Atto-647N. These are: DE=0.05, Lk=0.08, γ=0.85.

A plot of the stoichiometry versus the ALEX-2CDE filter was used to determine the required upper threshold that removes donor-only (S=1) and acceptor-only (S=0) molecules. In general, only molecules within the range S = 0.25–0.75 were considered in the final analysis. Doubly labeled photobleached molecules were further eliminated using the ALEX-2CDE (<14) and |TDX-TAA| (<0.5) filters as described before by Tomov et al.*(50)* and Kudryavtsev et al.*(24)*, respectively. These stringent filters guarantee elimination of unwanted signal, as described before *(24)*.

Lifetime was calculated using PAM after correction for instrument response factor (IRF). For the whole dataset, a double exponential decay function was used for the donor channel whereas a single exponential decay function was used for the acceptor channel *(49)*. Static and dynamic FRET lines were generated using PAM following previously published methods *(24, 25, 30)* and using an apparent linker length of 5 Å.

Subpopulation specific fluorescence lifetime analysis was also performed using PAM. However, for this type of analysis, the region of interest was first selected in the BurstBrowser module. Data were then systematically fit with one, two and three exponential functions to identify the best fit. Decisions were made based on analysis of the weighted residuals. Donor only and acceptor only species were also selected to verify that, in contrast to the FRET interval belonging to the open ensemble, they satisfactory fit a single exponential decay.

Theoretical FRET values were obtained by coarse-grained simulations using the FRET-restrained positioning and screening (FPS) software *(51)*. The dye dimensions were estimated to be 7.8, 4.5, and 1.5 Å for Atto-550 and 7.15, 4.5 and 1.5 Å for Atto-647N after minimization of their chemical structure using Maestro (Schrödinger). The linker lengths and widths used were 18 and 4.5 Å for both dyes. After performing accessible volume (AV) simulations, the corresponding mean transfer efficiency was calculated by assuming rapid fluctuations of the interdye distance occurring on time scales similar to the fluorescence lifetime of the donor (∼3.4 ns in the absence of the acceptor). This assumption is justified based on the anisotropy values reported in **Table 1** obtained for the labeled proteins.

### Dynamic Photon Distribution Analysis (PDA)

PDA analysis was performed using the PDAfit module built in PAM. Proximity histograms were reconstructed by binning the same dataset at 0.25, 0.5, 0.75, and 1 ms. Histogram library with a grid resolution for E=100 and a minimum number of photons of 10 per bin were chosen. The datasets were then fit using a dynamic three-state model. Distances calculated from lifetime analysis were fixed. The width of the distance distribution was also fixed at sigma=0.045. This was determined from the measurement of several static double-stranded doubly labeled DNA **(Figure S6)**. To assess robustness of the fit, PDA was repeated by systematically varying the initial value of the rate constants to 1, 0.5 and 0.75 ms^-1^ (min 0, max 10) while keeping the other settings identical. The results in **Table 4** represent the average of these three independent determinations.

### Species Selected Filtered FCS (fFCS)

Data were collected using the same setup described before in which we added two SPAD detectors (Excelitas Technologies) for a total of two parallel and two perpendicular detectors and increased the laser power to ∼30 μW. Briefly, the light was simultaneously split (50:50) and rotate by a polarizing cube. While the configuration of the perpendicular path is described before, the configuration of the parallel path is the following: dichroic filter ZT633rdc-UF1 (Chroma), donor emission filter ET585/65m (Chroma), and acceptor emission filter ET700/75 (Chroma). Fine-tuning of the system was performed such as very similar fluorescent intensity values (5% difference) were obtained for the two sets of detectors. FRET efficiency histograms and values of lifetime obtained for the 2- and 4-SPADs setups were identical.

Species selected filtered-Fluorescence Correlation Spectroscopy analysis was done using BurstBrowser module from PAM software. Microtime patterns for O_1_ and O_2_ states were obtained using FRET efficiency thresholds around the mean lifetime values obtained for O_1_ (E=0.17 to 0.25) and O_2_ (E=0.65 to 0.75) states. Signals from selected FRET efficiency region were cross-correlated after generating the appropriate TCSPC filters for the parallel and perpendicular channels. This eliminates the dead-time of TCSPC hardware and SPAD detectors. Four correlations functions, two auto correlation functions and two cross correlations functions between the species O_1_ and O_2_ were generated. The four curves were globally fitted using a single-component diffusion and single exponential kinetic term, as described by Felekyan et al. *(36)* and letting the amplitude for cross-correlation assume negative values. fFCS fit was carried out in FCSfit module from PAM. The diffusion time was fixed to 469 μs. This value was obtained from independent FCS measurements of reduced and oxidized PDI molecules at nanomolar concentrations. The ratio of the axial and lateral size of the confocal volumes were globally fixed, *ρ*=4.6. This was obtained from independent FCS measurements using singly labeled calibration samples such as double strand DNA or singly labeled PDI molecules at nanomolar concentrations. Capabilities of fFCS was tested using Holliday Junctions, as described elsewhere *(36)*.

### Size exclusion chromatography analyses

100 μL (100 μg) of a solution of PDI unbound and treated with 100 μM 16F16 or 100 μM PACMA-31 after reduction with DTT for 90 minutes at room temperature were loaded into a Superdex 200 HR 10/300 (Cytiva, USA) at a flow rate of 0.5 ml/min that was equilibrated with in Tris 20 mM, 150 mM NaCl at pH 7.4, 5 mM EDTA. Absorbance was monitored at 280 nm using an ÄKTApurifier system (Cytiva, USA).

## Data availability

All data are contained in the manuscript. PTU files are made available upon reasonable request by contacting the authors.

## Acknowledgments

We are thankful to Dr. Heyduk for granting access to the fluorimeter and for helpful discussions. We are also thankful to Dr. Frieden for granting access to the CD Spectrophotometer.

## Funding and additional information

This work was supported in part by grants R01 HL150146 (NP), R35 HL135775 (RF), and U01 HL143365 (RF) from the National Heart, Lung and Blood Institute.

## Authorship Contributions

M.C., R.F and N.P. designed the research; M.C. performed the research; M.C. and N.P. analyzed the data; M.C., R.F. and N.P. drafted the early version of the manuscript; N.P. wrote the final version of the manuscript; M.C., R.F. and N.P. edited the manuscript. All authors reviewed the manuscript.

## Conflict of interest

The authors declare that they have no conflicts of interest with the contents of this article.

## Supplementary Materials

**Figure S1.**
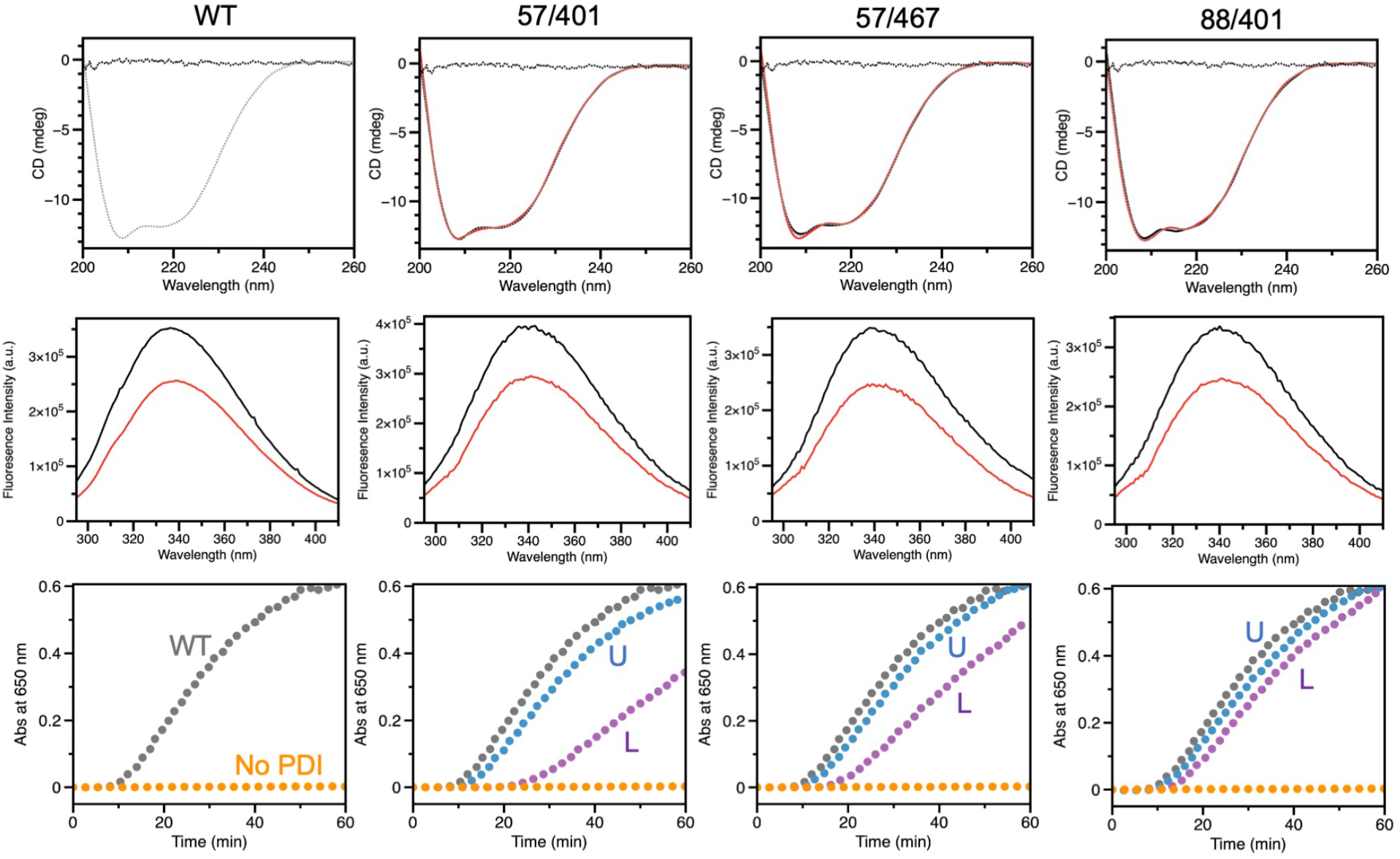
Structural and functional characterization of the FRET variants. Shown are far-UV CD spectra (top row), response to GSSG (red) and GSH (black) monitored by intrinsic fluorescence (center row) and progress curves for insulin reductase activity assay (bottom row) of PDI wild-type and variants. The labels U and L indicate unlabeled and doubly labeled protein samples, respectively. Results for the variant 88/467 are reported in **Figure 1** of the manuscript.

**Figure S2.**
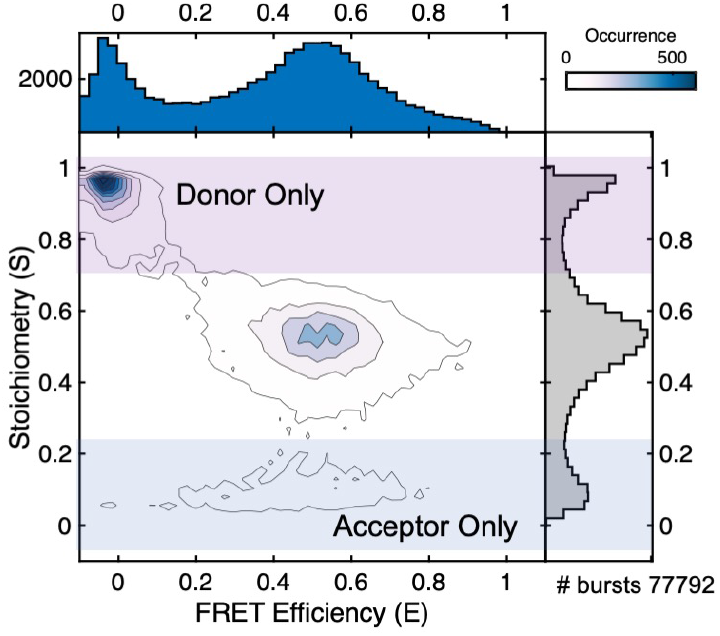
2D FRET efficiency vs Stoichiometry plot of reduced PDI 88/467 before cleanup. Highlighted are Donor only (S>0.75, magenta) and Acceptor only (S<0.25, blue) populations. These species were discarded as they are not relevant to our analysis. On average, doubly labeled species account for 18-25% of the total.

**Figure S3.**
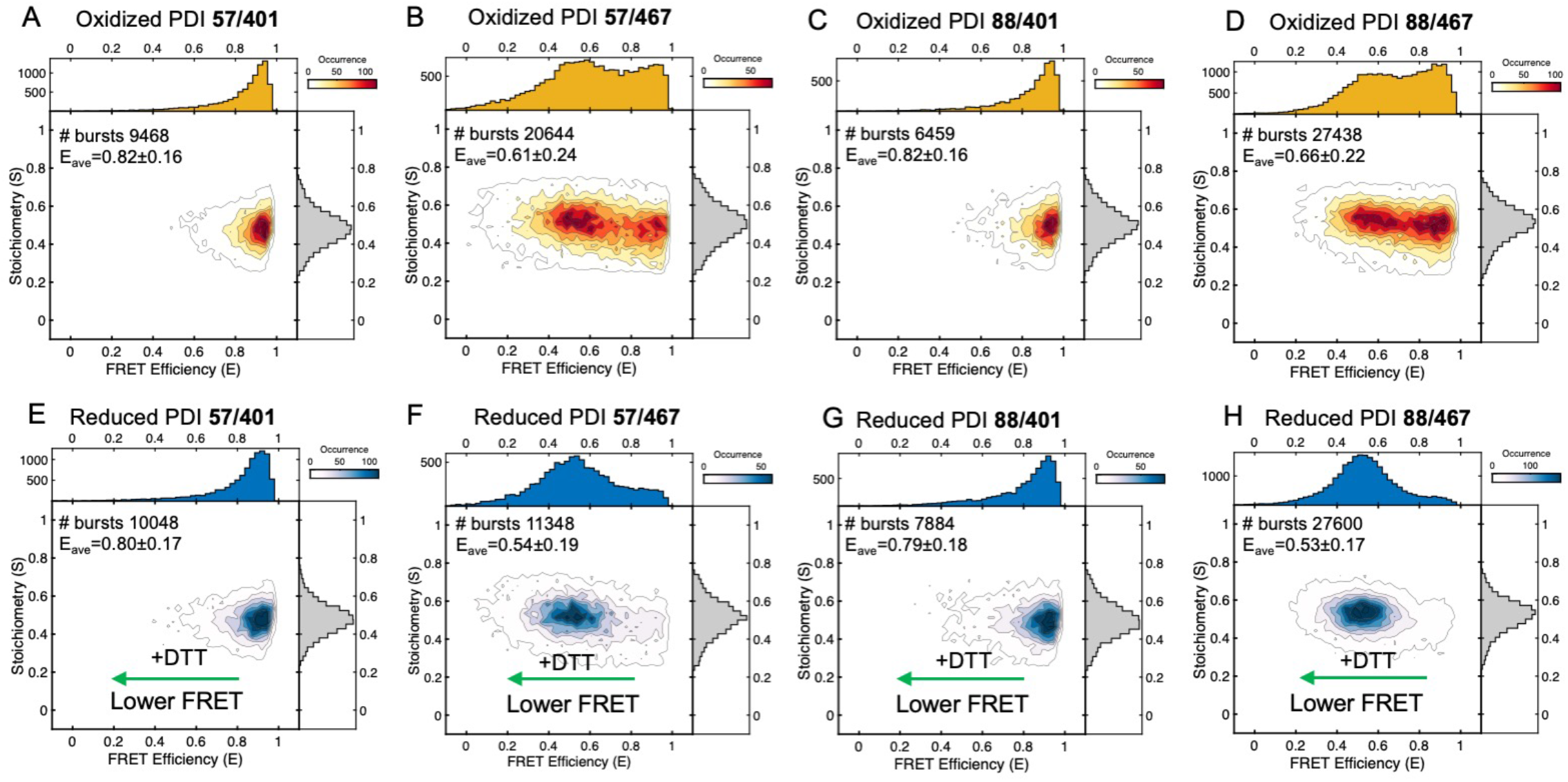
2D FRET efficiency vs Stoichiometry plots of PDI 57/401, PDI 57/467, PDI 88/401 and PDI 88/467 under non-reducing (top, yellow) and reducing conditions (bottom, blue). Note how addition of DTT, while shifting E_ave_ towards lower FRET, does not affect stoichiometry.

**Figure S4.**
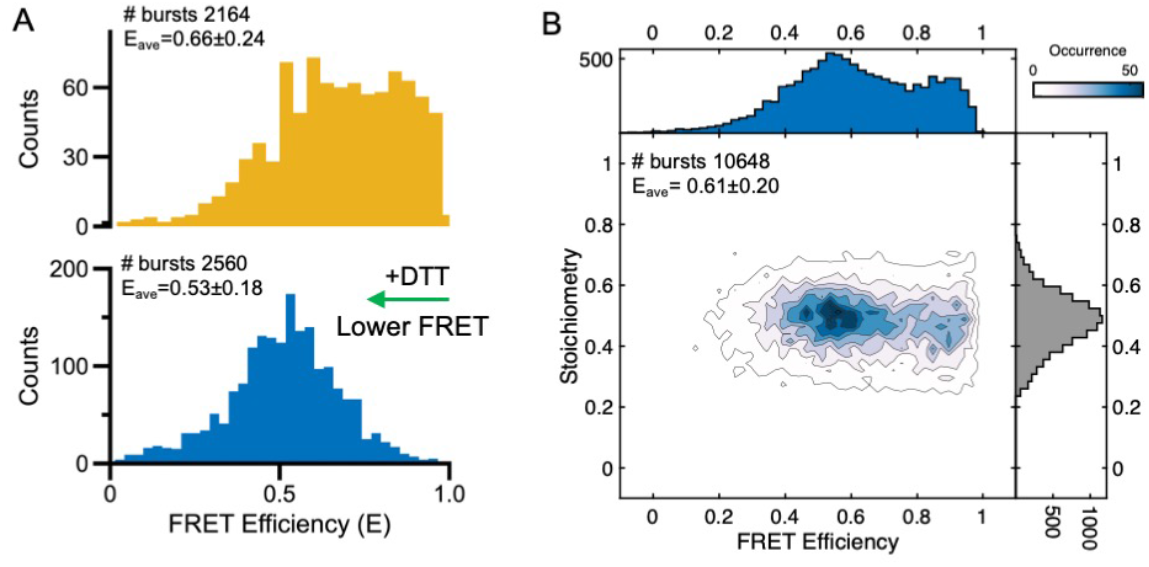
Control experiments to characterize PDI 88/467. **(A)** PDI 88/467 labeled with sulfo-Cy3/sulfo-Cy5 azide (Lumiprobe) under non reducing (top, yellow) and reducing (bottom, blue) conditions. PDI 88/467 labeled with Cy dyes displays similar high to low FRET transition in the presence of DTT compared to PDI 88/467 labeled with Atto dyes. **(B)** PDI 88/467 labeled with Atto dyes in the presence of 1 mM GSH. GSH is less potent than DTT when used at the same concentration.

**Figure S5.**
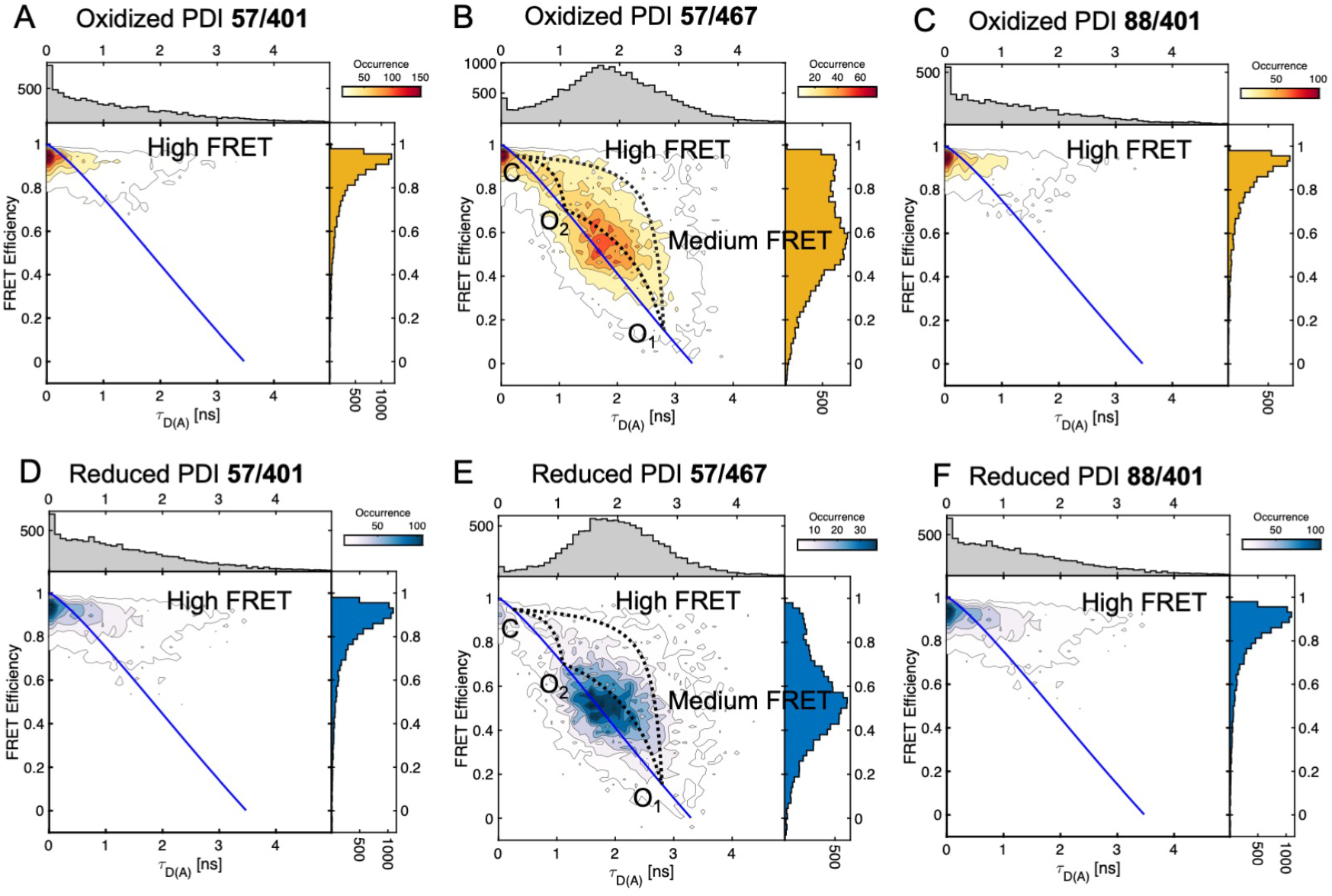
2D FRET efficiency vs lifetime plots of PDI 57/401, PDI 57/467 and PDI 88/401 under non-reducing (top, yellow) and reducing (1 mM DTT) conditions (bottom, blue). Static FRET lines (solid blue lines) are shown in each plot. Due to high FRET, PDI 57/401 **(A** and **D)** and PDI 88/401 **(B** and **E)** show only a hint of dynamics, which manifests as a small but significant deviation of the high FRET ensemble towards the right of the static FRET line. Also evident in these plots is the shift toward lower FRET induced by DTT. PDI 57/467 **(C** and **F)**, in contrast to PDI 57/401 and PDI 88/401, but similar to PDI 88/467 (**Figure 3** of the main text), shows a very clear dynamic signature documenting dynamic exchange between closed (high FRET, C) and open (medium FRET, O) ensembles. These two ensembles are characterized by mean fluorescence lifetime values of ∼0.25 and ∼1.8 ns, respectively. The open ensemble of PDI 57/467, similar to PDI 88/467, is shifted toward the right of the static FRET line indicating fast dynamics between open states. Using the same methodology described in the main text, we identified O_1_ (1.1±0.2 ns) and O_2_ (2.8±0.3 ns), which are shown in the plot along the dynamic FRET lines (black dotted lines) that connect them.

**Figure S6.**
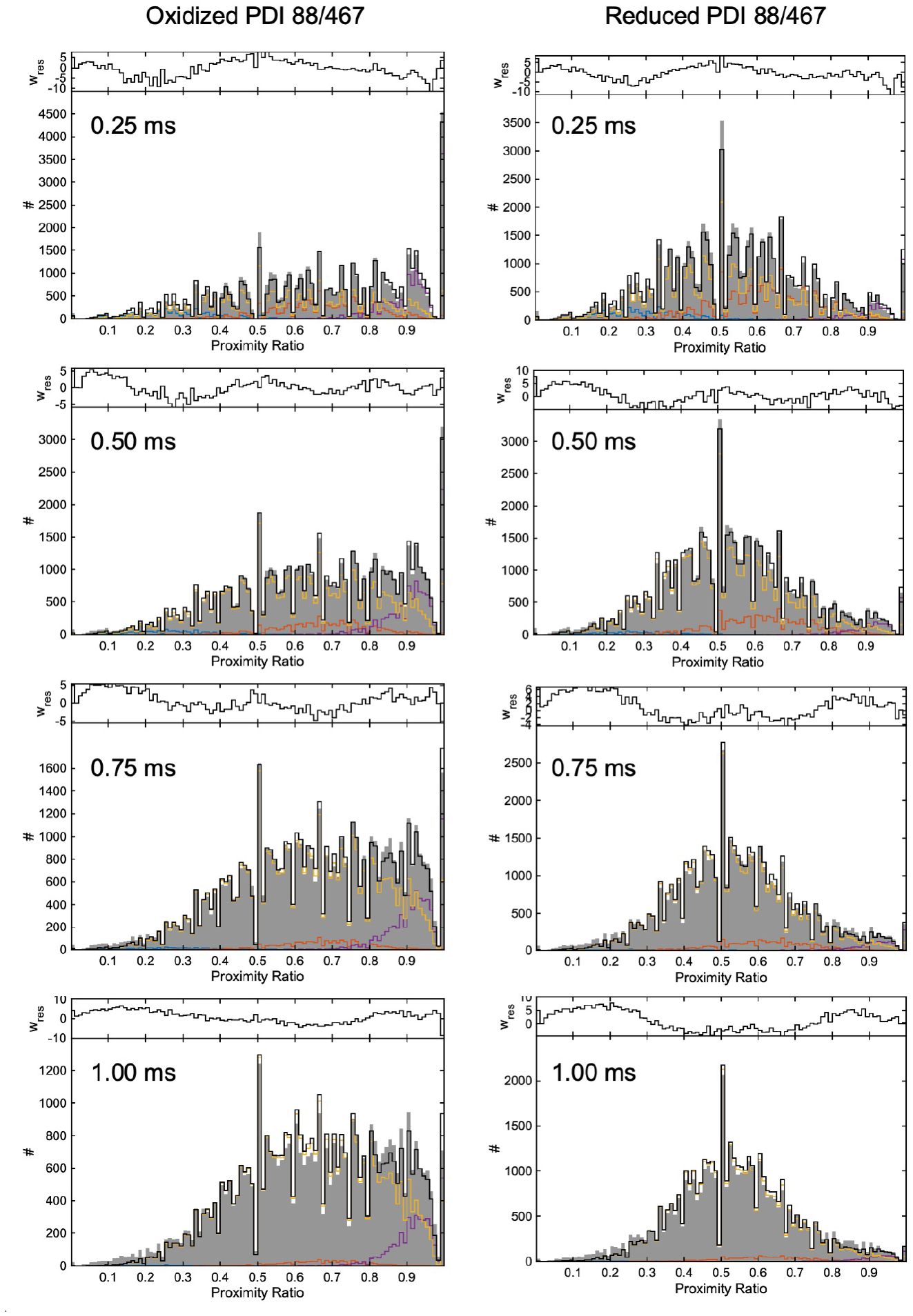
PDA analysis of PDI 88/467. PDA was performed on dataset binned at 0.25, 0.5, 0.75 and 1 ms. Photons from each burst were used to build a proximity ratio (PR) histogram. The resulting histogram was then fitted using a Monte Carlo approach for simulating the burst-wise histogram using a dynamic three-state model. To assess robustness of the fit, PDA was repeated by systematically varying the initial value of the rate constants to 1, 0.5 and 0.75 ms^-1^ (min 0, max 10) while keeping the other settings identical. Corresponding weighted residuals are shown above each plot.

**Figure S7.**
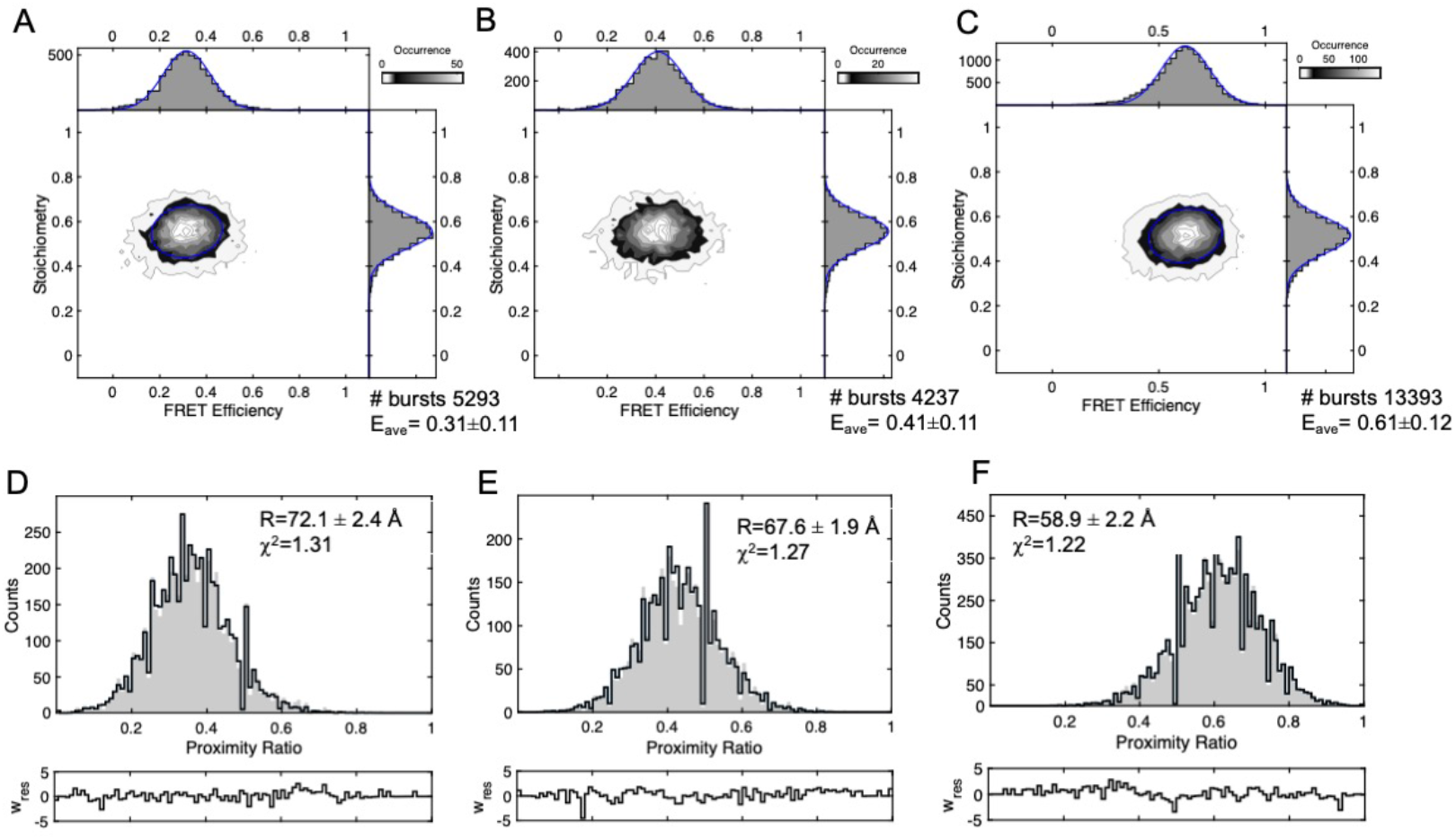
Analysis of static double-stranded DNA constructs. 2D plots **(A, B and C)** and PDA **(D, E and F)** analysis of DNA duplexes with probes separated by 19 **(A)**, 17 **(B)** and 14 **(C)** base pairs. Single-stranded DNA molecules were purchased (IDT Inc., Coralville, LA) and fluorescent dyes (Atto550/647N) were attached to amino dT residues obtained by substituting T to iAmMC6T. dsDNA molecules were formed by hybridization. Experimental conditions are 100 pM in TBS-Tween 0.01%. FRET histograms best fit to a one Gaussian distribution (blue). Note how the standard deviation for static species is significantly smaller compared to values obtained in this work for PDI **(Table 1)**, supporting the view that PDI adopts multiple conformations in solution. PDA was performed on dataset binned at 1 ms. Photons from each burst were used to build a proximity ratio histogram. The resulting histogram was then fitted using a Monte Carlo approach for simulating the burst-wise histogram using one Gaussian. Corresponding weighted residuals are shown below each plot.

**Figure S8.**
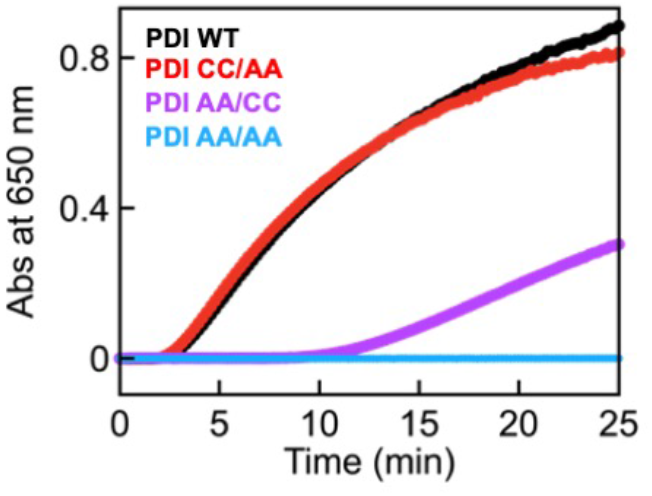
Reductase activity of the PDI 88/467 and active site mutants. Reductase activity of PDI 88/467 (WT, black) and active site variants PDI 88/467 C53A/C56A/C397AC400A (AA/AA, blue), PDI 88/467 C397AC400A (CC/AA, red) and PDI C53A/C56A (AA/CC, magenta) monitored by the insulin assay. Note how the catalytic activity of PDI CC/AA is similar to PDI WT but different from PDI AA/CC, whose catalytic activity is compromised. Among the two active sites, the one in the **a** domain is the most important for insulin reduction. PDI AA/AA is catalytically inactive, as expected, since no longer contains cysteine residues in the active sites.

**Figure S9.**
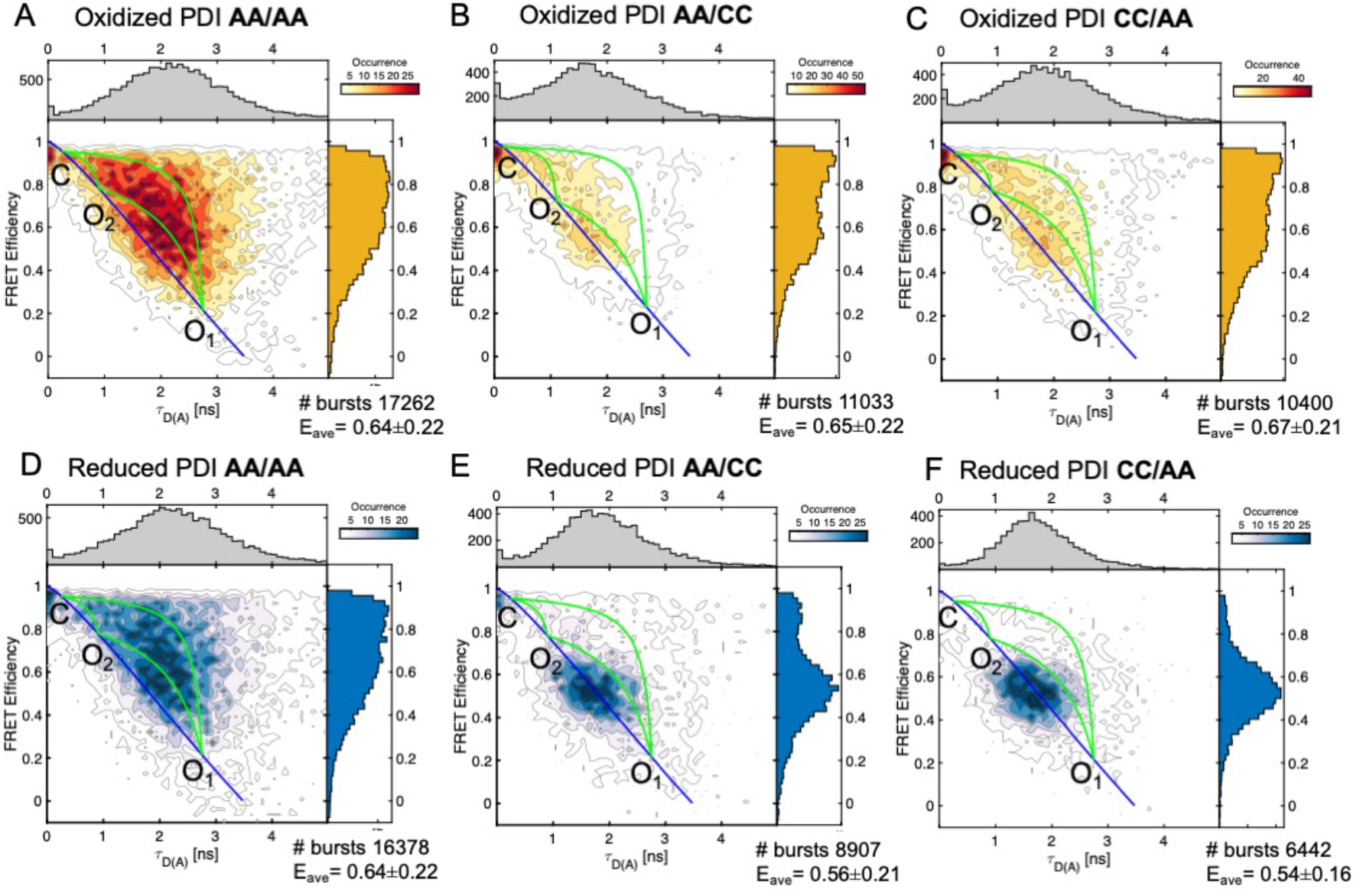
2D FRET efficiency vs lifetime plots of active site variants under non-reducing (top, yellow) and reducing conditions (bottom, blue). Shown are static (solid blue lines) and dynamic (solid green lines) FRET lines connecting the FRET states. The lines were drawn as described in the main text. The position of C, O_1_ and O_2_ is indicated. The fraction of each population was obtained by PDA and is reported in **Table 4**.

## Notes

### Competing Interest Statement

The authors have declared no competing interest.

### Summary of Updates

New title

## References

1. F. Hatahet, L. W. Ruddock, Protein disulfide isomerase: a critical evaluation of its function in disulfide bond formation. Antioxid Redox Signal 11, 2807–2850 (2009).

2. M. Matsusaki et al., The Protein Disulfide Isomerase Family: from proteostasis to pathogenesis. Biochim Biophys Acta Gen Subj, (2019).

3. L. Ellgaard, L. W. Ruddock, The human protein disulphide isomerase family: substrate interactions and functional properties. EMBO Rep 6, 28–32 (2005).

4. Y. Wu, D. W. Essex, Vascular thiol isomerases in thrombosis: The yin and yang. J Thromb Haemost, (2020).

5. L. Wang, J. Yu, C. C. Wang, Protein disulfide isomerase is regulated in multiple ways: Consequences for conformation, activities, and pathophysiological functions. Bioessays, e2000147 (2020).

6. P. V. S. Oliveira et al., Protein disulfide isomerase plasma levels in healthy humans reveal proteomic signatures involved in contrasting endothelial phenotypes. Redox Biol 22, 101142 (2019).

7. M. Matsusaki et al., The Protein Disulfide Isomerase Family: from proteostasis to pathogenesis. Biochim Biophys Acta Gen Subj 1864, 129338 (2020).

8. R. Flaumenhaft, B. Furie, Vascular thiol isomerases. Blood 128, 893–901 (2016).

9. J. Chiu, P. J. Hogg, Allosteric disulfides: Sophisticated molecular structures enabling flexible protein regulation. J Biol Chem 294, 2949–2960 (2019).

10. J. Ahamed et al., Disulfide isomerization switches tissue factor from coagulation to cell signaling. Proc Natl Acad Sci U S A 103, 13932–13937 (2006).

11. S. Kumar et al., An allosteric redox switch in domain V of beta2-glycoprotein I controls membrane binding and anti-domain I autoantibody recognition. J Biol Chem 297, 100890 (2021).

12. J. Cho et al., Protein disulfide isomerase capture during thrombus formation in vivo depends on the presence of beta3 integrins. Blood 120, 647–655 (2012).

13. J. Li et al., Platelet Protein Disulfide Isomerase Promotes Glycoprotein Ibalpha-Mediated Platelet-Neutrophil Interactions Under Thromboinflammatory Conditions. Circulation 139, 1300–1319 (2019).

14. S. Chakravarthi, C. E. Jessop, N. J. Bulleid, The role of glutathione in disulphide bond formation and endoplasmic-reticulum-generated oxidative stress. EMBO Rep 7, 271–275 (2006).

15. G. Tian, S. Xiang, R. Noiva, W. J. Lennarz, H. Schindelin, The crystal structure of yeast protein disulfide isomerase suggests cooperativity between its active sites. Cell 124, 61–73 (2006).

16. C. Wang et al., Structural insights into the redox-regulated dynamic conformations of human protein disulfide isomerase. Antioxid Redox Signal 19, 36–45 (2013).

17. L. Peng, M. I. Rasmussen, A. Chailyan, G. Houen, P. Hojrup, Probing the structure of human protein disulfide isomerase by chemical cross-linking combined with mass spectrometry. J Proteomics 108, 1–16 (2014).

18. R. A. Romer et al., The flexibility and dynamics of protein disulfide isomerase. Proteins 84, 1776–1785 (2016).

19. R. B. Freedman et al., ‘Something in the way she moves’: The functional significance of flexibility in the multiple roles of protein disulfide isomerase (PDI). Biochim Biophys Acta Proteins Proteom 1865, 1383–1394 (2017).

20. M. Okumura et al., Dynamic assembly of protein disulfide isomerase in catalysis of oxidative folding. Nat Chem Biol 15, 499–509 (2019).

21. E. I. Biterova et al., The crystal structure of human microsomal triglyceride transfer protein. Proc Natl Acad Sci U S A 116, 17251–17260 (2019).

22. M. Chinnaraj et al., Bioorthogonal Chemistry Enables Single-Molecule FRET Measurements of Catalytically Active Protein Disulfide Isomerase. Chembiochem, (2020).

23. H. Sanabria et al., Resolving dynamics and function of transient states in single enzyme molecules. Nat Commun 11, 1231 (2020).

24. V. Kudryavtsev et al., Combining MFD and PIE for accurate single-pair Forster resonance energy transfer measurements. Chemphyschem 13, 1060–1078 (2012).

25. S. Kalinin, A. Valeri, M. Antonik, S. Felekyan, C. A. Seidel, Detection of structural dynamics by FRET: a photon distribution and fluorescence lifetime analysis of systems with multiple states. J Phys Chem B 114, 7983–7995 (2010).

26. C. Wang et al., Human protein-disulfide isomerase is a redox-regulated chaperone activated by oxidation of domain a’. J Biol Chem 287, 1139–1149 (2012).

27. I. Grossman et al., Single-molecule spectroscopy exposes hidden states in an enzymatic electron relay. Nat Commun 6, 8624 (2015).

28. J. J. Alston, A. Soranno, A. S. Holehouse, Integrating single-molecule spectroscopy and simulations for the study of intrinsically disordered proteins. Methods, (2021).

29. I. V. Gopich, A. Szabo, Theory of the energy transfer efficiency and fluorescence lifetime distribution in single-molecule FRET. Proc Natl Acad Sci U S A 109, 7747–7752 (2012).

30. A. Barth et al., Dynamic interactions of type I cohesin modules fine-tune the structure of the cellulosome of Clostridium thermocellum. Proc Natl Acad Sci U S A 115, E11274–E11283 (2018).

31. I. Grossman-Haham, G. Rosenblum, T. Namani, H. Hofmann, Slow domain reconfiguration causes power-law kinetics in a two-state enzyme. Proc Natl Acad Sci U S A 115, 513–518 (2018).

32. M. Okumura, K. Noi, K. Inaba, Visualization of structural dynamics of protein disulfide isomerase enzymes in catalysis of oxidative folding and reductive unfolding. Curr Opin Struct Biol 66, 49–57 (2020).

33. I. V. Gopich, A. Szabo, Single-molecule FRET with diffusion and conformational dynamics. J Phys Chem B 111, 12925–12932 (2007).

34. Y. Santoso, J. P. Torella, A. N. Kapanidis, Characterizing single-molecule FRET dynamics with probability distribution analysis. Chemphyschem 11, 2209–2219 (2010).

35. B. Hellenkamp et al., Precision and accuracy of single-molecule FRET measurements-a multi-laboratory benchmark study. Nat Methods 15, 669–676 (2018).

36. S. Felekyan, S. Kalinin, H. Sanabria, A. Valeri, C. A. Seidel, Filtered FCS: species auto- and cross-correlation functions highlight binding and dynamics in biomolecules. Chemphyschem 13, 1036–1053 (2012).

37. J. D. Stopa, K. M. Baker, S. P. Grover, R. Flaumenhaft, B. Furie, Kinetic-based trapping by intervening sequence variants of the active sites of protein-disulfide isomerase identifies platelet protein substrates. J Biol Chem 292, 9063–9074 (2017).

38. B. G. Hoffstrom et al., Inhibitors of protein disulfide isomerase suppress apoptosis induced by misfolded proteins. Nat Chem Biol 6, 900–906 (2010).

39. K. S. Cole et al., Characterization of an A-Site Selective Protein Disulfide Isomerase A1 Inhibitor. Biochemistry 57, 2035–2043 (2018).

40. S. Xu et al., Discovery of an orally active small-molecule irreversible inhibitor of protein disulfide isomerase for ovarian cancer treatment. Proc Natl Acad Sci U S A 109, 16348–16353 (2012).

41. H. Mazal et al., Tunable microsecond dynamics of an allosteric switch regulate the activity of a AAA+ disaggregation machine. Nat Commun 10, 1438 (2019).

42. C. Wang et al., Plasticity of human protein disulfide isomerase: evidence for mobility around the X-linker region and its functional significance. J Biol Chem 285, 26788–26797 (2010).

43. L. J. Byrne et al., Mapping of the ligand-binding site on the b’ domain of human PDI: interaction with peptide ligands and the x-linker region. Biochem J 423, 209–217 (2009).

44. R. H. Bekendam et al., A substrate-driven allosteric switch that enhances PDI catalytic activity. Nat Commun 7, 12579 (2016).

45. E. Ruben et al., The J-elongated conformation of beta2-glycoprotein I predominates in solution: implications for our understanding of antiphospholipid syndrome. J Biol Chem 295, 10794–10806 (2020).

46. N. Pozzi, D. Bystranowska, X. Zuo, E. Di Cera, Structural Architecture of Prothrombin in Solution Revealed by Single Molecule Spectroscopy. J Biol Chem 291, 18107–18116 (2016).

47. A. N. Kapanidis et al., Fluorescence-aided molecule sorting: analysis of structure and interactions by alternating-laser excitation of single molecules. Proc Natl Acad Sci U S A 101, 8936–8941 (2004).

48. A. C. Ferreon, C. R. Moran, Y. Gambin, A. A. Deniz, Single-molecule fluorescence studies of intrinsically disordered proteins. Methods Enzymol 472, 179–204 (2010).

49. W. Schrimpf, A. Barth, J. Hendrix, D. C. Lamb, PAM: A Framework for Integrated Analysis of Imaging, Single-Molecule, and Ensemble Fluorescence Data. Biophys J 114, 1518–1528 (2018).

50. T. E. Tomov et al., Disentangling subpopulations in single-molecule FRET and ALEX experiments with photon distribution analysis. Biophys J 102, 1163–1173 (2012).

51. S. Kalinin et al., A toolkit and benchmark study for FRET-restrained high-precision structural modeling. Nat Methods 9, 1218–1225 (2012).

